# A regulatory toolkit of arabinose-inducible artificial transcription factors for Gram-negative bacteria

**DOI:** 10.1101/2022.11.30.518220

**Authors:** Gita Naseri, Hannah Raasch, Emmanuelle Charpentier, Marc Erhardt

**Affiliations:** Institut für Biologie, Humboldt-Universität zu Berlin, Philippstrasse 13, 10115 Berlin, Germany; Max Planck Unit for the Science of Pathogens, Charitéplatz 1, 10117 Berlin, Germany

**Author notes:** **Corresponding authors**: Marc Erhardt,; Gita Naseri.

**Keywords:** Arabinose, Artificial transcription factor, *Escherichia coli*, Gram-negative bacteria, *Salmonella*, Synthetic biology

## Abstract

The Gram-negative bacteria *Salmonella* Typhimurium and *Escherichia coli* are important model organisms, powerful prokaryotic expression platforms for biotechnological applications, and pathogenic strains constitute major public health threats. To facilitate new approaches for research, biomedicine, and biotechnological applications, we developed a set of arabinose-inducible artificial transcription factors (ATFs) using CRISPR/dCas9 and *Arabidopsis-derived* DNA-binding proteins, allowing to control gene expression in *E. coli* and *Salmonella* over a wide inducer concentration range. As a proof-of-concept, we employed the developed ATFs to engineer a *Salmonella* biosen<u>sor</u> strain, SALSOR 0.2 (SALmonella biosenSOR 0.2), which responds to the presence of alkaloid drugs with quantifiable fluorescent output. We demonstrated that SALSOR 0.2 was able to detect the presence of the antitussive noscapine alkaloid with ~2.3-fold increased fluorescent signal over background noise compared to a previously described biosensor. Moreover, we used plant-derived ATFs to control β-carotene biosynthesis in *E. coli*, which resulted in ~1.6-fold higher β-carotene production compared to expression of the biosynthesis pathway using a strong constitutive promoter. The arabinose-inducible ATFs reported here thus enhance the synthetic biology repertoire of transcriptional regulatory modules that allow tuning protein expression in the Gram-negative model organisms *Salmonella* and *E. coli*.

## Introduction

Gram-negative bacteria, including *Escherichia coli* and *Salmonella enterica*, have been genetically engineered for high-titer production and secretion of difficult-to-express proteins such as conotoxins, antimicrobial peptides, and a malaria vaccine via the type-III secretion systems (T3SS)^1, 2, 3^, and have also been harnessed in cancer therapy for delivering drugs to the host cells^4, 5, 6, 7^ as they can grow in aerobic or anaerobic conditions in solid tumors^8, 9, 10^. In addition, pathogenic strains of *E. coli* and *S. enterica* are among the world’s most significant public health problems due to increasing antibiotics resistance^11^. Through their T3SSs, they inject effector proteins directly into host cells^12, 13^. However, the complex mechanisms of the underlying gene regulatory networks (GRNs) controlling expression of many important virulence factors is poorly understood^14, 15, 16^. To understand such regulatory networks, for example, during infection or to reprogram cellular behaviors^17^, it is highly desirable to have robust genetic tools such as orthogonal transcriptional regulators that allow to artificially control these networks^18^. Toward this goal, Davis *et al*. (2011) have reported the development of four synthetic promoters *PproA, B, C*, and *D*, to constitutively modulate the expression of endogenous or heterologous proteins in *E. coli*^19^. Later, Cooper *et al* (2017) applied the *Ppro* series of promoters to improve the tunability of protein expression in *Salmonella^20^*. However, employing constitutive elements to modulate the gene expression may result in a metabolic burden on the cell because the expression of heterologous proteins competes with other cellular processes and may be undesirable when an orthogonal (minimal interference with the native cellular processes) and controllable (allowing expression at the desired time) system is needed^21^. To address the aforementioned challenges, artificial transcription factors (ATFs) allowing for temporal, tight, and tunable gene expression are favorable. Some ATFs have been developed for synthetic biology applications in both prokaryotic and eukaryotic organisms^22, 23, 24, 25, 26, 27^. However, only a limited number of ATFs are available for Gram-negative bacteria^24, 25, 26, 27^, and to our knowledge, no dedicated ATF to control gene expression in *Salmonella* has been reported.

Few inducible ATFs, based on catalytically-inactive CRISPR-associated protein Cas9 (dCas9) and Cas12a (dCas12) DNA binding domains (DBDs) have been developed *e.g*., in *E. coli*^24, 25, 26, 27^ and *Paenibacillus polymyxa*^28^. To activate the bacterial endogenous transcriptional machinery, these ATFs are equipped with activation domains (ADs), e.g. the omega subunit of RNA polymerase (RNAP)^26^ or bacterial enhancer-binding proteins (bEBPs)^27^. However, these CRISPR/dCas-derived ATF systems require a specific genetic background that harbors deletions of the omega subunit and bEBPs, which may lead to fitness defects and might be unfavorable as the required genomic manipulation is restricted to specific host backgrounds. To overcome these issues, Dong *et al*. (2018) used the AD of the bacterial transcription factor SoxS to activate gene expression in *E. coli*^24^.

Here, we used the same approach as described by Dong *et al*. (2018) to establish ATFs using dCas9 DBD and SoxS AD for tunable gene expression in *Salmonella*. We further evaluated a novel class of ATFs with heterologous DBDs based on plant-specific transcription factors (TFs) from *Arabidopsis thaliana*^22, 29^, combined with the SoxS AD^23^ in order to extend our library of ATFs that allow for inducible gene expression in Gram-negative bacteria. The expression of ATFs is typically controlled by exogenous inducers. Unlike *E. coli*^30, 31^, little attention has been paid to modulating the expression level derived from the arabinose-inducible *araBAD* promoter (*P_BAD_*) in *Salmonella*. Because the arabinose induction system is slightly different in *Salmonella*, compared to *E. coli*^32, 33^, we first genetically modified the L-arabinose (hereafter, arabinose) catabolic pathway in *Salmonella enterica* serovar Typhimurium resulting in ~4.5-to ~6.5-fold improved induction of *P_BAD_*-derived gene expression. For our collection of arabinose-inducible ATFs, we further defined an optimal ‘arabinose induction window’ (0.01% to 0.1% arabinose) in *Salmonella*, that resulted in a high heterologous gene expression level with only minimal effects on bacterial growth. Finally, as a proof-of-concept for the diverse applications of ATFs, we engineered a *Salmonella* strain to function as a sensitive biosensor for alkaloid drugs and an *E. coli* strain as a microbial cell factory to produce β-carotene.

## RESULTS

### Design of arabinose-inducible artificial transcription factors

A fundamental aim of our work was to develop inducible, heterologous regulators that allow to genetically reprogram gene regulatory networks of the Gram-negative bacteria *Salmonella* and *E. coli*. For this purpose, we generated arabinose-inducible ATFs derived from widely different DBDs: CRISPR/dCas9 and plant heterologous TFs (**Fig. 1**). In order to evaluate the performances of these ATFs to induce gene expression, we developed a set of reporter and expression plasmids. Chromosomally or plasmid-derived expressed ATFs were placed under the control of *P_BAD_*. In the absence of arabinose, dimeric AraC acts as a repressor, where one monomer binds to the operator *O_2_*, and another monomer binds to the *I_1_* half-site in *P_BAD_*, which results in the formation of a DNA loop that prevents RNAP from binding to *P_BAD_*. In the presence of arabinose and upon binding of arabinose to AraC, binding of the AraC-arabinose complex to the *I_2_* half-site in *P_BAD_* is allosterically induced, while binding to *O_2_* is decreased. In this configuration, AraC acts as activator, promoting the binding of RNAP to *P_BAD_*, which activates expression of the ATF. Next, the ATF targets its BS(s) located upstream of a weak synthetic promoter in a way that the AD contacts RNAP to promote its binding to the promoter. As a result the expression of the target gene is activated^34^.

**Fig. 1.**
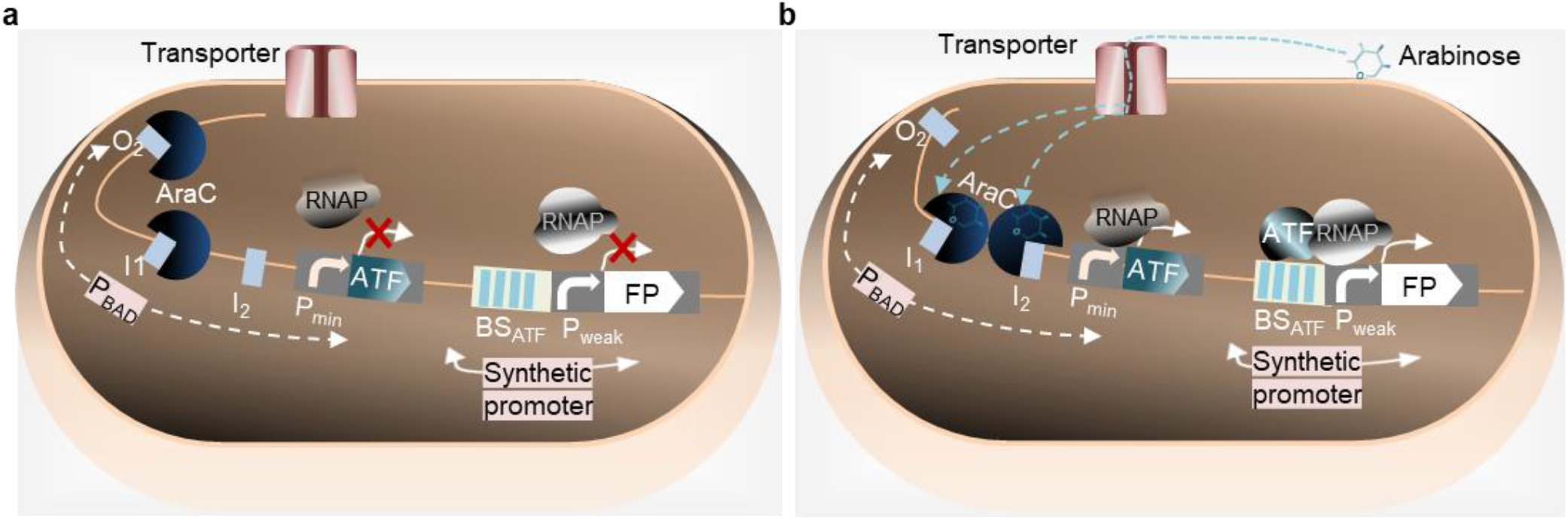
Principle of arabinose-inducible ATFs established in this study. **a** Inducer-OFF state: In the absence of arabinose, AraC dimer binds to *O_2_* and *I_1_* half-sites, causing DNA looping, which prevents RNAP from accessing the promoter. As a result, the ATF is not expressed. Therefore, the expression of FP that is controlled by BS of ATF is prevented. **b** Inducer-ON state: Following the addition of arabinose, it enters the cell by a transporter. Arabinose-bound AraC dimer changes configuration and binds *I_1_* and *I_2_* half-sites of *P_BAD_*, activating the transcription of the ATF. The ATF targets its BS within a synthetic promoter, controlling FP expression. The interaction of the ATF and RNAP leads to increased FP transcription from the synthetic promoter compared to the OFF state^34^. To simplify the figure, the RBSs and terminators located, respectively, upstream and downstream of ATF and FP are not shown. Abbreviations: ATF, artificial transcription factor; BS, binding site; FP, fluorescent protein; *I_1_*, *I_2_* and *O_2_* represent DNA binding half-sites. *GOI*, gene of interest; *P_BAD_*, arabinose-inducible *araBAD* promoter; *P_min_*, minimal synthetic promoter containing the −35 and −10 regions of the essential elements; *P_weak_*, weak promoter; RNAP, RNA polymerase.

### Development of an arabinose-based toolkit for a wide range of transcriptional outputs in *Salmonella*

To maximize inducible gene expression from *P_BAD_*, it is necessary to identify conditions that shift the equilibrium from arabinose catabolism to maximal induction of gene expression while having a minimum effect on cellular fitness^10, 31^. The *P_BAD_* promoter has been extensively used to express heterologous and endogenous genes in *E. coli* and *Salmonella*. While the regulatory mechanism of the arabinose induction system has been well-characterized in *E. coli*^30, 31, 35^, few studies have been performed on characterizing the arabinose induction system in *Salmonella*^30, 36^. In contrast to *E. coli*, *araE* is the only gene that encodes for an arabinose-specific transporter in *Salmonella*^32, 36^. In the native system, intracellular arabinose induces expression of the *araBAD* operon (which encodes three enzymes AraB, AraA, and AraD that convert arabinose to d-xylulose-5-phosphate to provide carbon and energy for cellular metabolisms) and of the *araE* gene (**Fig. 2a**)^10^.

**Fig. 2.**
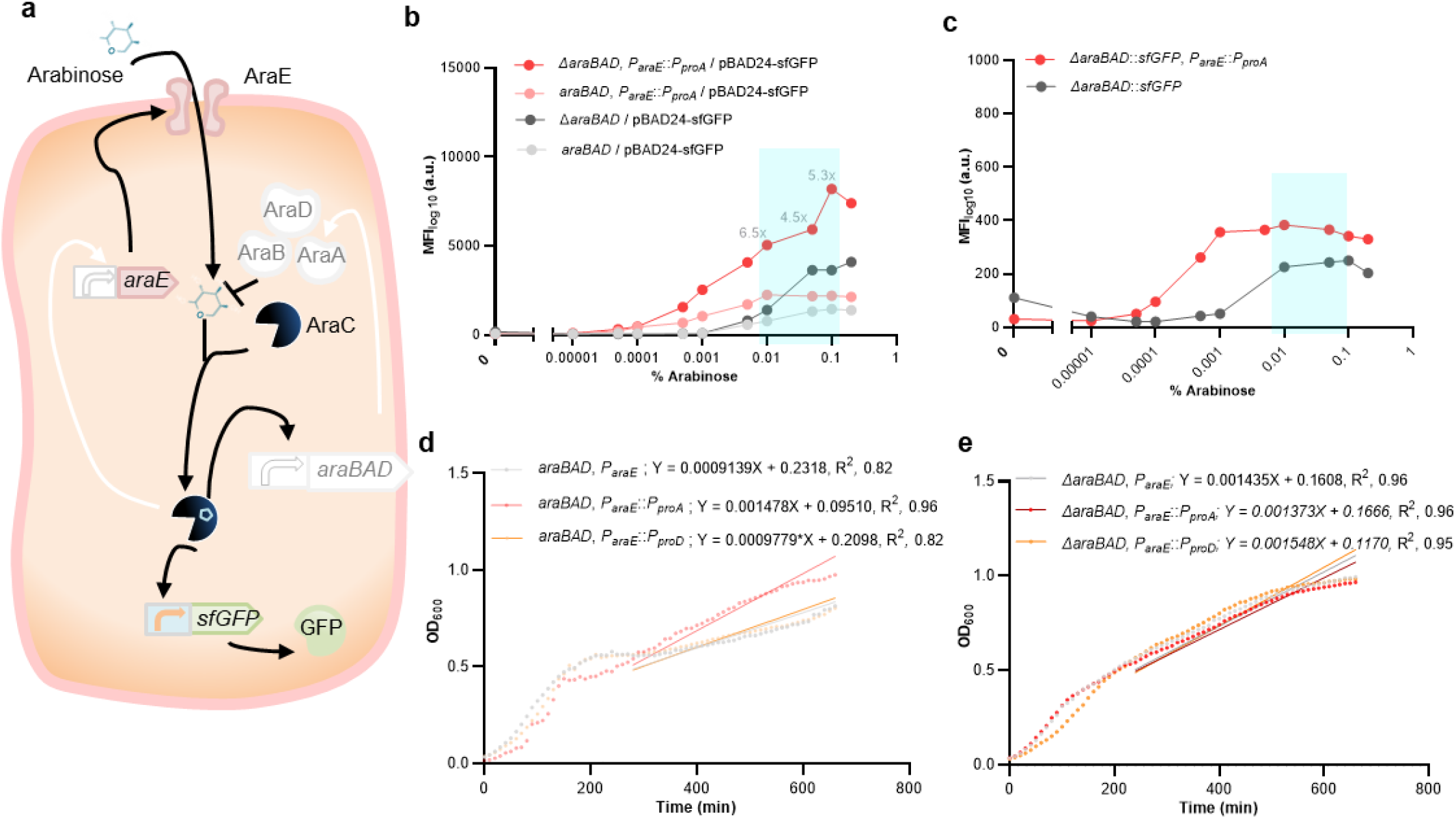
Optimization of the arabinose-based toolkit in *Salmonella*. **a**. Regulatory network of the native arabinose utilization system in *S*. Typhimurium^32^, including the reporter cassette used in this study. The system consists of *araE*, encoding the arabinose transporter AraE essential for arabinose uptake, the *araBAD* operon, encoding enzymes for arabinose metabolism, and the regulator AraC-encoding gene. High amounts of intracellular arabinose activate *araC* expression, which stimulates expression from the promoters *P_BAD_* and *P_araE_* (in the absence of arabinose, AraC represses *P_BAD_*-derived expression). Indicated in white shading are the mutants characterized in this study: deletion of the *araBAD* operon and constitutive *araE* expression at different levels. As a reporter, either plasmid-or chromosomal-based GFP expression system under the control of *P_BAD_* was used. To simplify the figure, the possible negative feedback of AraC is not shown, as it mainly contributes to provide a constant AraC level^37^. Characterizing dose-dependency of arabinose-based gene expression from plasmid (**b**) or chromosome (**c**) in *S*. Typhimurium LT2. Light grey, wild-type; dark grey, *araBAD* deletion mutant; light red, mutant with decoupled arabinose-dependent transporter-reporter system; dark red, mutant with *araBAD* deletion and decoupled arabinose-dependent transporter-reporter system. As a reporter, sfGFP under the control of the *P_BAD_* promoter, encoded on a plasmid (pBAD24-sfGFP) or in the chromosome (*ΔaraBAD::sfGFP*), was used. sfGFP-expression was measured by flow cytometry in presence of 0.00001%, 0.00005%, 0.0001%, 0.0005%, 0.001%, 0.005%, 0.01%, 0.05%, 0.1%, and 0.2% arabinose. The ‘induction window’ is highlighted in blue. Growth-curves of strains mutated for (**d**) arabinose-independent expression of the transporter and (**e**) arabinose-independent expression of transporter and deleted arabinose catabolism system in the presence of 0.2% arabinose in *S*. Typhimurium LT2 background. The growth of cells expressing *araE* from the weak *P_proA_* (light red)^20^ or strong *PproD* (light orange)^20^ constitutive promoters was compared with that of cells with arabinose-dependent (araE) expression from the native *P_araE_* in cells with or without deleted *araBAD* operon. Data are the average of three biological replicates. Abbreviations: a.u., arbitrary units; MFI, mean fluorescent intensity; *P_BAD_*, arabinose-responsive promoter; OD600, optical intensity at 600 nm; R^2^, the statistical measure of regression predictions in Goodness of Fit test; sfGFP, super-fold green fluorescent protein; X and Y, parameters of the linear regression equation. The full data are shown in **Data S1**.

Like in *E. coli*, where addition of 0.2% arabinose results in maximal induction *P_BAD_*, 0.2% arabinose is also commonly used to induce both chromosomal and episomal *P_BAD_* in *S*. Typhimurium^10^. In order to investigate the induction kinetics of the arabinose system in *Salmonella*, we quantified the output of a *P_BAD_*-dependent fluorescent protein reporter after induction during mid-log growth phase with various concentrations of arabinose ranging from 0.00001% to 0.2% using flow cytometry (**Fig. 2b** and **Supplementary Fig. 1**). Our data demonstrated that addition of 0.0005% arabinose was sufficient to maximally induce *P_BAD_*-controlled expression of plasmid-encoded GFP and 0.05% arabinose for the chromosomal construct (**Fig. 2b**, light grey). Additionally, deleting the *araBAD* operon, so that arabinose cannot be metabolized, resulted in remarkedly increased reporter protein expression compared to the *araBAD^+^* strain (**Fig. 2b**, dark grey). A decoupled arabinose transporter-reporter system, where *araE* was constitutively expressed using the relatively weak *P_proA_* promoter^20^, resulted in maximal induction at a 10-fold lower arabinose concentration than the *araE*^+^ strain, where *araE* is under control of its native (arabinose-inducible) promoter *P_araE_* (**Fig. 2b**, light grey and light red, 0.01% and 0.1%). A combination of decoupled arabinose transporter-reporter system and deletion of the *araBAD* operon resulted in improved *P_BAD_*-dependent reporter gene expression over a wide range of 0.0005% to 0.2% arabinose (**Fig. 2b** and **Fig. 2c**, dark red).

A growth curve analysis further showed that, in presence of 0.2% arabinose, *P_proA_*-controlled expression of *araE* (P*_proA_*-*araE*) resulted in slower growth during early exponential growth in *araBAD^+^* background (**Fig. 2d**, red), compared to native, arabinose-dependent *araE* expression in the wild-type strain (*P_araE_-araE*) (**Fig. 2d**, grey). Upon entering later growth phases (from time point 240 min), the *P_proA_-araE* strain grew better compared to the *P_araE_-araE* strain. In contrast, strong constitutive *araE* expression from the *P_proD_* promoter resulted in similar growth compared to native arabinose-responsive *araE*-expression in later growth phases. These results suggest that constitutive arabinose-independent, low-level expression of AraE is optimal for an arabinose-inducible expression system. In 0.05% or lower arabinose concentrations, the growth of strains expressing *araE* from either native, *P_proA_* or *PproD* was similar (**Supplementary Fig. 2a, b**, and **c**). In high arabinose concentrations (0.1% and 0.2%), the reduced growth of strains expressing *araE* from either native, *P_proA_* or *PproD* is compensated upon *araBAD* operon deletion (**Fig. 2e, d**, and **Supplementary Fig. 2d, e**, and **f**). The high activity of the *P_BAD_*-derived reporter system (**Fig. 2b** and **c**, highlighted in blue) with minimal growth defect (**Supplementary Fig. 2**, blue curves) implies an optimal ‘arabinose induction window’ ranging from arabinose concentrations of 0.01% to 0.1%.

### Arabinose-inducible CRISPR/dCas9-derived ATF library

We next established an arabinose-inducible ATF system, derived from a previously developed CRISPR/dCas9-derived ATF to activate gene expression in *E. coli*, employing a catalytically inactive version of *Streptococcus pyogenes* Cas9 (dCas9) as DBD^24^. The dCas9 binds to scRNA targeting the *J1*-derived synthetic promoter. The scRNA is fused to a MS2 hairpin sequence, which recruits a MS2 coating protein (MCP)-fused to a R93A mutant of the SoxS activator domain, which has previously been shown to effectively activate transcription^24, 28^. The SoxS activator domain is recruited to dCas9 DBD by a scaffold RNA (scRNA) extended with a hairpin MS2 sequence. We, here, employed the codon-optimized dCas9 and SoxS (R93A)^24^ system to establish an arabinose-inducible CRISPR/dCas9-derived ATF library in *Salmonella* (**Fig. 3a**).

**Fig. 3.**
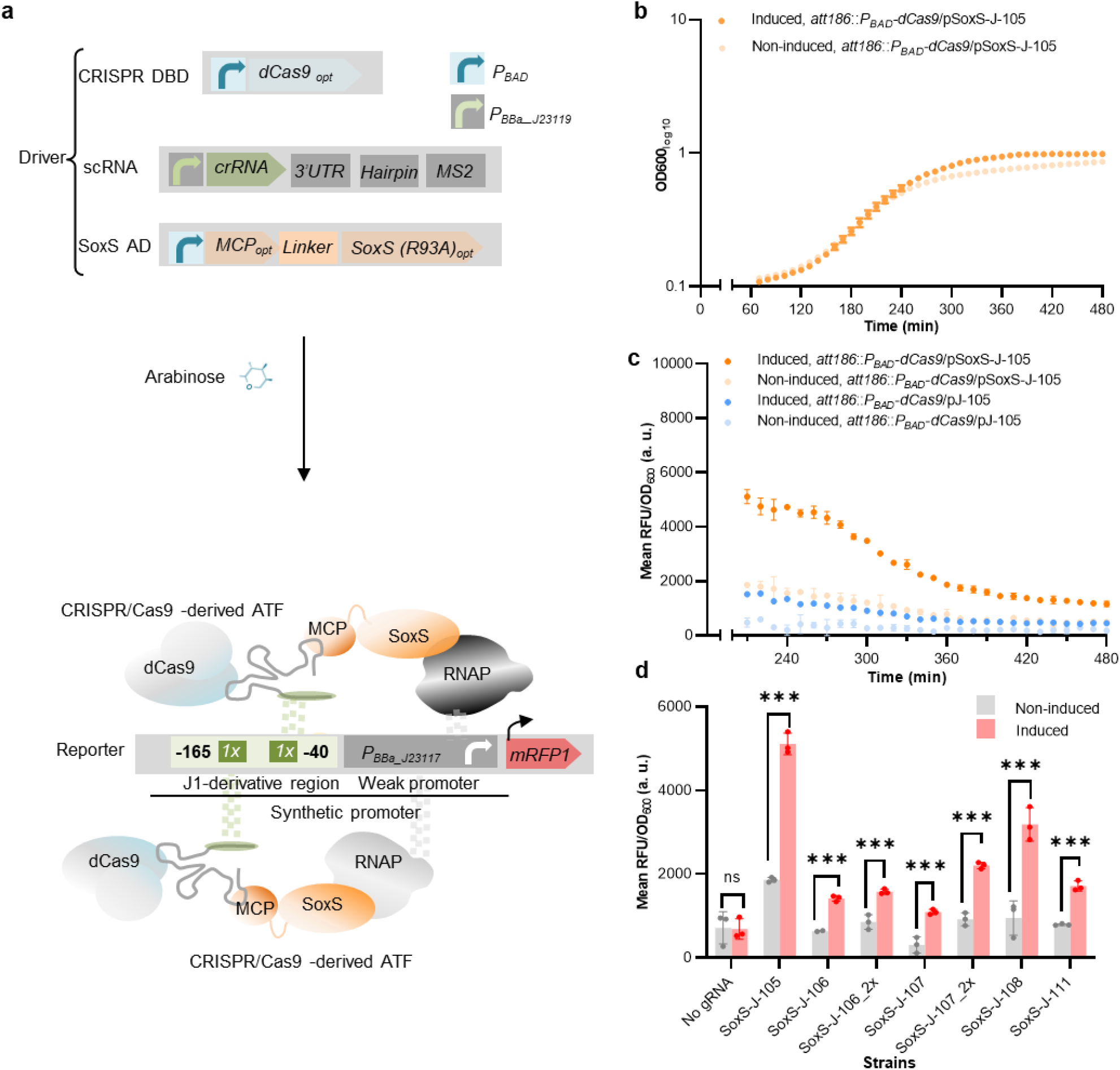
Arabinose-inducible CRISPR/dCas9-derived ATF library for tunable gene expression in *Salmonella*. **a** Schematics showing CRISPR/dCas9-derived ATFs developed in the present study. The driver cassette (top) of CRISPR/dCas9-derived ATFs comprises i) DBD cassette containing a *P_BAD_* at the 5’ end and *S. pyogenes* dCas9 (codon-optimized for expression in *S*. Typhimurium LT2 (dCas9 opt)), ii) scRNA cassette, encoding the crRNA and MS2 RNA hairpins under control of the constitutive *P_BBa-J23119_*^38^, and iii) AD cassette encoding MCP fused to SoxS(R93A) AD (codon-optimized for expression in *S*. Typhimurium LT2) via 5aa-linker under control of PBAD. In the presence of arabinose, dCas9 and SoxS are expressed. SoxS AD is recruited by MCP to the dCas9-scRNA complex via the MS2 RNA hairpin. The expressed driver targets a *J1*-derived synthetic promoter placed upstream of the gene of interest in the reporter cassette (bottom) via (a sequence in the) scRNA. The *J1*-derived synthetic promoter has potential crRNA target sites upstream of a weak P_BBa_J23117_^24^. The schematics shows the condition in which the *J1*-derived synthetic promoter harbors two copies of ATF BS, located between −165 and −40 (to TSS). The interaction of the ATF and RNAP induces the transcription of *mRFP1^34^*. Growth curve (**b**) and time course of mRFP1 expression level (**c**) in cells expressing the CRISPR/dCas9-derived ATF targeting J105 in *S*. Typhimurium LT2 background. Cells harbor chromosomally-integrated dCas9, plasmid-encoded MCP-linker-<u>S</u>oxS(R93A), J<u>105</u>-targeting scRNA and *<u>J</u>1*-derived synthetic promoter to control mRFP1 expression. Light color, non-inducing medium (arabinose^-^); dark color, inducing medium (arabinose^+^). The statistically significant difference in expression for cells expressing CRISPR/dCas9-derived ATF targeting J105 under inducing medium (dark orange) compared to non-inducing medium (light orange) is shown in **Data S2**. **d** Library of arabinose-inducible, CRISPR/Cas9-derived ATFs in *Salmonella*. Fluorescence output was measured in the non-inducing medium (grey) and inducing medium (red). Data are expressed as the mean ± SD of the RFU obtained from three biological replicates, normalized to the OD_600_. Asterisks indicate a statistically significant difference from the non-inducing medium (two-sided *t-*test; *p ≤ 0.05, \*\*\**p* ≤ 0.001). For induction, 0.05% arabinose was used. Abbreviations: aa, amino acid; AD, activation domain; a. u., arbitrary units; ATF, artificial transcription factor; BS, binding site; DBD, DNA binding domain; dCas9, catalytically inactive Cas9; gRNA, guide RNA; MCP, MS2 coating protein; mRFP1, monomeric red fluorescent protein 1; OD_600_, optical density at 600 nm; RFU, relative fluorescence units; scRNA, scaffold RNA; RNAP, RNA polymerase. The full data are shown in **Data S2**.

Dong *et al*. (2018)^24^ engineered an ATF system, in which dCas9 and scRNA expression are under the control of *P_BAD_* and SoxS is expressed from a constitutive promoter. The scRNAs target the regions J105, J106, J107, J108, and J111 within the *J1*-derived synthetic promoter, resulting in high transcriptional output^24^. We adapted this system and used *P_BAD_* to control the expression of the relatively large dCas9 protein and SoxS AD in *S*. Typhimurium to minimize the negative effect of CRISPR/Cas-derived ATF expression on growth (**Fig. 3b**, dark orange). In contrast, the “scRNA” cassette was expressed from the strong *PBBa-J23119* promoter^38^. Additionally, dCas9 was integrated into the chromosome at the attachment site of coliphage 186 (*att186*)^39^. Expression of the CRISPR/Cas-derived ATF targeting region J105^24^ by addition of 0.05% arabinose did not result in a growth defect of *S*. Typhimurium (**Fig. 3b**). Our data show that *P_BAD_*-controlled expression of the SoxS AD is required for the CRISPR/dCas9-derived ATF to activate reporter gene expression (**Fig. 3c**, dCas9 without SoxS AD under inducing conditions (dark blue) and non-inducing conditions (light blue); dCas9 and SoxS AD under inducing conditions (dark orange) and non-inducing conditions (light orange)). Although we observed leaky reporter gene expression in the absence of the inducer, likely due to leaky expression from *P_BAD_* (light orange), adding 0.05% arabinose resulted in significantly enhanced reporter gene expression (dark orange). However, expression of the reporter gene decreased dramatically and reached the background level at later growth phases (after 480 min) (**Fig. 3c**, dark orange). Therefore, we performed arabinose-inducible experiments to characterize our ATFs in mid-log phase cultures.

We selected the scRNA targeting regions J105, J106, J107, J108, and J111 for further characterization. Moreover, to further extend the size of our library, we designed *J1* derivatives harboring two copies of J106 and J107 situated in a narrow region that is required for effective gene activation (see also **Fig. 1** and **Fig. 3d**, J106_2x compared to J106, and J-107_2x compared to J-107). Our driver cassette resulted in the inducible expression of CRISPR/dCas9-derived ATFs controlling expression of the fluorescent reporter protein in *Salmonella* (**Fig. 3d**). In *E. coli* (**Supplementary Fig. 3a**), we were not successful in obtaining similar results for the ATFs targeting J106 and J107 as reported by Dong *et al*. (2018), suggesting that gene activation is likely sensitive to the established expression system in *E. coli*, which is consistent with prior results^24^. Further examination of the total population distribution showed that reporter gene expression was induced in less than 27% of *E. coli* cells (**Supplementary Fig. 3b**) and in more than 63% of *Salmonella* cells (**Supplementary Fig. 4**) in the case of strong CRISPR/Cas-derived ATF under the tested condition (*E. coli* 0.2% arabinose, and *Salmonella* 0.05% arabinose), suggesting that an inadequate amount of arabinose was used. This result highlights that developing ATFs for inducible gene expression in bacteria is challenging. However, the CRISPR/Cas9-derived ATF described here is a valuable tool for tunable gene expression in *Salmonella*.

### Arabinose-inducible plant-derived ATF library

An ideal regulatory toolkit should enable bioengineers to linearly induce gene expression over a wide dynamic range. Therefore, we aimed to extend the library of arabinose-inducible ATFs for bioengineering applications in *Salmonella* and *E. coli*. Inspired by MacDonald *et al*. (2021)^40^ who demonstrated that the eukaryotic transcription factor QF from the fungus *Neurospora crassa* can be used in *E. coli* to activate transcription, we developed a new class of ATFs using heterologous TFs derived from plant for tunable gene expression in Gram-negative bacteria. Over 1000 plant-specific TFs have been identified in higher plants, grouped into diverse families according to conserved motifs in their DBD^41^.

In order to establish plant-derived ATFs for bacterial gene expression, full-length plant TFs GRF9, ANAC102 and JUB1-derived ATFs (which function as weak, medium and strong ATFs in yeast^22, 23, 29^), and the DBD of JUB1, JUB1_DBD_ (which functions as medium ATF in yeast) were used. The plant TFs were fused to the bacterial SoxS(R93A) activation domain^24, 28^ optimized for expression in *Salmonella* (**Fig. 4a**). We first evaluated the capacity of chromosomal or plasmid-based expression of the plant-derived ATFs under *P_BAD_*-control to activate transcription in *E. coli*. Following the addition of inducer (0.2% arabinose), ATFs are expressed from the driver cassette (**Fig. 4a**), and target two copies of their BSs within a synthetic promoter resulting in expression of the reporter sfGFP. We show that an ATF based on the full-length plant JUB1 TF, rather than only on its DBD, displays the strongest capacity to activate gene expression in *E. coli* (**Fig. 4b**) consistent with our previous findings of the transcription activation capacity of ATFs in yeast^23^.

**Fig. 4.**
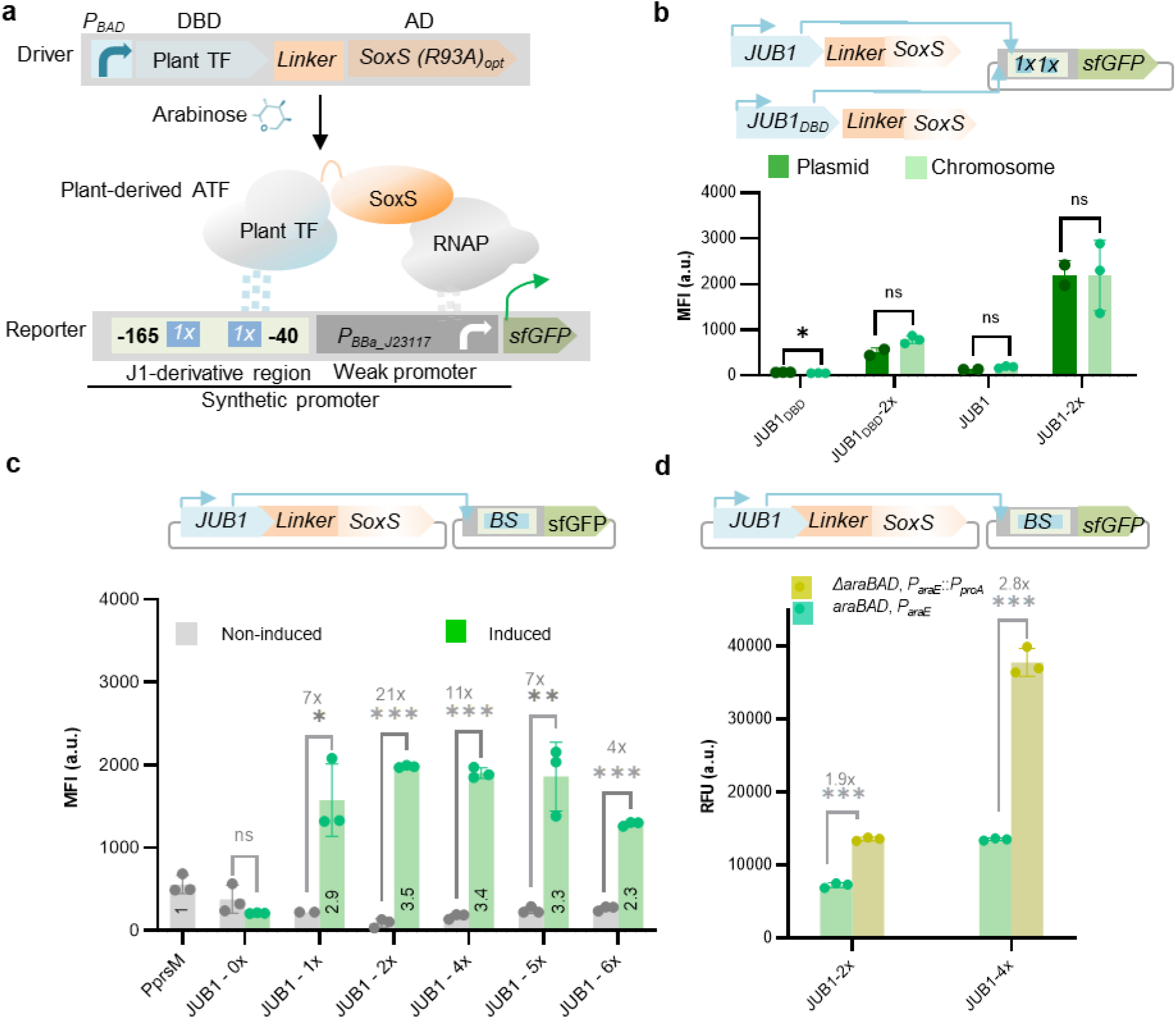
Arabinose-inducible, plant-derived ATFs for tunable gene expression in Gram-negative bacteria. **a** Schematics showing plant-derived ATFs developed in the present study. The driver cassette contains a *P_BAD_* at the 5’ end controlling expression of a plant TF fused via 5 aa-linker to SoxS(R93A) activation domain. In the presence of arabinose, the expressed ATF binds a *J1*-derived synthetic promoter upstream of *P_BBa_J23117_* promoter^24^. In the schematics a reporter cassette with a synthetic promoter harboring two copies of TF-BS located between −165 and −40 (to TSS) is shown as example. Interaction between the ATF and RNAP drives reporter sfGFP expression^34^. **b** Arabinose-inducible, JUB1-derived ATFs encoded on a plasmid or in the chromosome in *E. coli* Full-length plant JUB1 or only its DBD was chromosomally integrated at the *att186* site (dark green), or expression plasmids harboring full-length plant JUB1 or only its DBD were introduced into cell by transformation (light green). The ATF transactivation capacity in combination with two copies of BS was tested in inducing medium. The strains harboring driver cassette, but no reporter cassette were used as a negative control. **c** Arabinose-inducible, JUB1-derived ATF in *Salmonella*. The transactivation capacity of JUB1-derived ATFs in combination with the one, two, four, five or six copies of BS was tested in wild-type background. Grey, non-inducing medium; green, inducing medium; Constitutive promoter *P_prsM_*, positive control; JUB1-0x, negative control. **d** Optimization of arabinose-inducible, JUB1-derived ATFs toolkit in *Salmonella*. The transactivation capacity of the JUB1-derived ATF in combination with two or four copies of BS was tested in strain with deleted *araBAD* and constitutively expressing *araE* (from *P_proA_*) (see **Fig. 2**) (dark yellow) and wild-type background (green) in inducing medium. Data are expressed as the mean ± SD of the MFI (**b**, **c**) or RFU (**d**) obtained from three biological replicates. Asterisks indicate a statistically significant difference (*t*-test; ns, not significant; \**p* ≤ 0.05; \*\**p* ≤ 0.01; \*\*\**p* ≤ 0.001). To simplify the figure, ribosome binding site and terminator are not shown. For induction, 0.2% and 0.05% arabinose were used in *E. coli* and *S*. Typhimurium LT2, respectively. Abbreviations: aa, amino acid; ATF, artificial transcription factor; a. u., arbitrary units; BS, binding site; JUB1, JUNGBRUNNEN1; JUB1-0X, synthetic promoter without binding site of JUB1-derived ATF; MFI, mean fluorescence intensity; RNAP, RNA polymerase; RFU, relative fluorescent unit; sfGFP, super-fold green fluorescent protein; TF, transcription factor. The full data are shown in **Data S3**.

Since we observed similar transcriptional outputs for both plasmid- and chromosomally-derived expression systems, we implemented plasmid-based expression of the driver cassette in the following experiments as it may be favorable in some cases over genomic integration due to their easy manipulation. The JUB1-derived ATF, in combination with one, two, four, five and six copies of its BS was then characterized in *S*. Typhimurium. The obtained driver/reporter displayed an inducible range of more than ~21-fold when induced with 0.05% arabinose (**Fig. 4c**). Of note, all of the ATFs resulted in higher sfGFP reporter output than the control strain expressing sfGFP from the constitutive *P_proA_* promoter (**Fig. 4c**, up to a ~3.5-fold, number inside the columns). All tested combinations of GRF9- or ANAC102-derived ATFs and their BSs resulted in low transcriptional outputs in *Salmonella* (~0.4-to ~0.6-fold of that observed for the control strain, **Supplementary Fig. 5**) despite being categorized as medium and strong regulators in yeast^23^. To further establish a dynamic arabinose-inducible system for precise temporal gene expression control in bacteria, we next characterized the JUB1-derived ATFs, together with two or four copies of the BS, in a S. Typhimurium background deleted for the *araBAD* operon and harboring a decoupled arabinose transporter-reporter system (**Fig. 2b**, *ΔaraBAD, P_proA_::P_proE_*). The developed inducible transcription system displayed high expression strength when induced with 0.05% arabinose (~2.8-fold, **Fig. 4d**,).

### Plant-derived ATF capacity to establish *Salmonella* as biosensor for alkaloid drug discovery

A key obstacle in the microbial production of high-value chemicals is to identify enzymes that can improve yield^42^. d’Oelsnitz *et al*. (2022) developed a genetically encoded biosensor based on the multidrug-resistance regulator RamR repressor from *S*. Typhimurium that enables detection of the benzylisoquinoline alkaloid (BIA) group of plant therapeutic alkaloids^43^. They engineered *E. coli* for heterologous expression of RamR and demonstrated the utility of this sensor as a tool to detect BIAs, including the anti-tumor alkaloid noscapine (NOS)^44^ by incorporating its binding site into a synthetic promoter driving expression of a fluorescent reporter gene. *Salmonella* strain expressing genomic RamR under the control of its native promoter was engineered to harbor a synthetic promoter with the RamR BS driving sfGFP expression, termed SALSOR 0.1 (**Fig. 6a**). Moreover, we engineered the SALSOR 0.2 strain, which, in addition to harboring the synthetic RamR-sfGFP reporter module, expresses chromosomally RamR encoded under the control of our above-developed arabinose-inducible JUB1-derived ATF in combination with synthetic promoter harboring four copies of JUB1 BS (**Fig. 6b**). As shown in **Fig. 6c**, the SALSOR 0.1 biosensor strain was able to detect NOS. Low background signals (in the absence of NOS) generally lead to an increased signal-to-noise ratio and thus a better detection (lower detection limit) of the ligand. Therefore, we reasoned that employing the arabinose-inducible JUB1-derived ATF would allow us to increase the expression levels of the RamR repressor. As expected, when using the SALSOR 0.2 biosensor, the background signal was reduced by 30% compared to the SALSOR 0.1 biosensor (**Fig. 6c**).

**Fig. 5.**
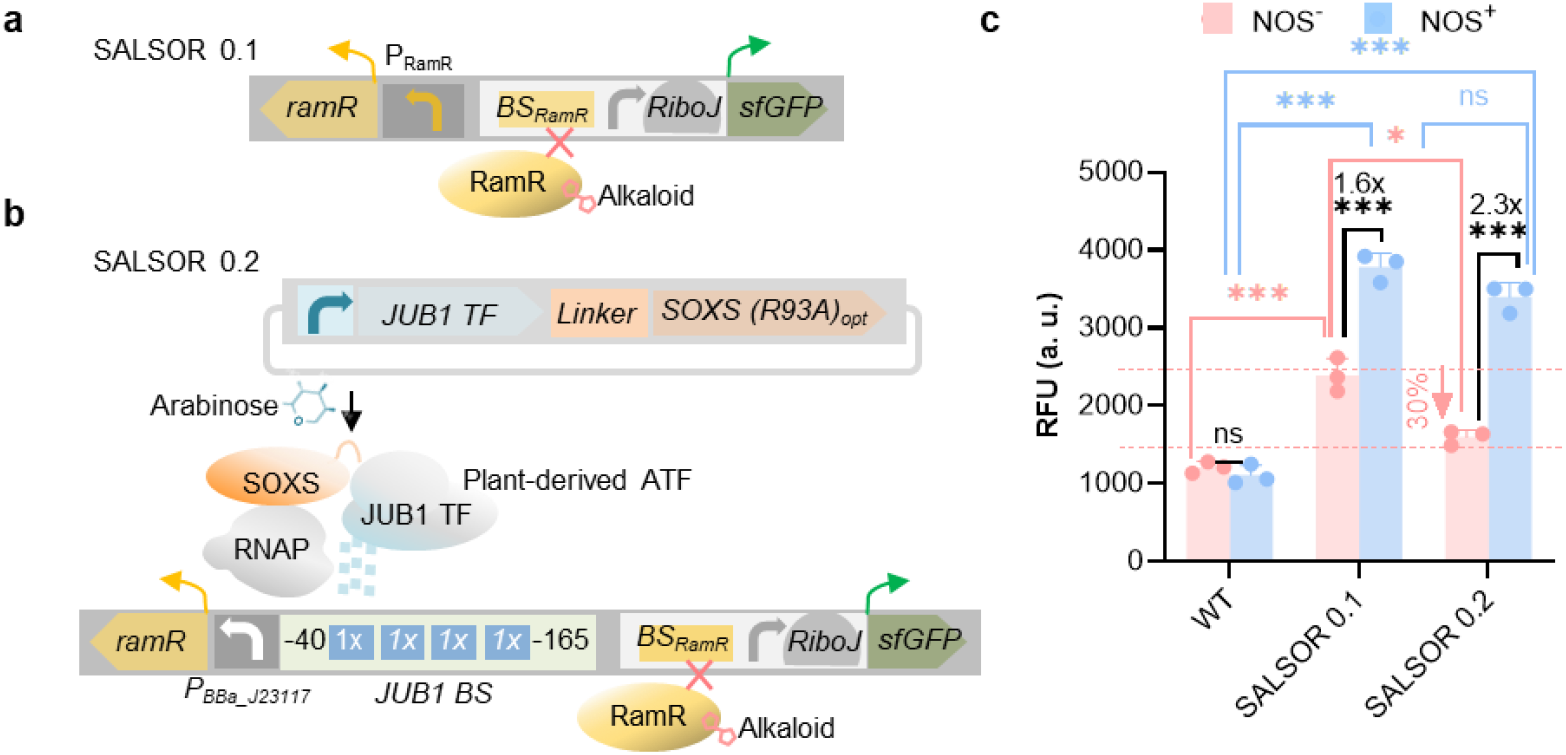
*Salmonella* biosensor responsive to alkaloid. **a** Schematic of the genetic circuit used in SALSOR 0.1 strain. The RamR sensor’s native constitutive promoter (present in *S*. Typhimurium LT2) was implemented to express RamR. Expressed RamR targets a synthetic promoter consisting of RamR sensor’s operator BS_RamR_ and UP element, followed by RiboJ-RBS (75 nucleotide sequence consisting of a satellite RNA of tobacco ringspot virus derived ribozyme followed by a 23-nucleotide hairpin immediately downstream to help expose the RBS)^43, 45^ that controls sfGFP expression. In the presence of NOS, RamR interaction with BSRamR is inhibited. **b** Schematic of the genetic circuit used in SALSOR 0.2 strain. Plasmid-encoded JUB1-derived ATF is expressed in the presence of 0.05% arabinose. A synthetic promoter containing four copies of JUB1 BS fused to weak P_BBa_J23117_^24^ was used to express RamR sensor. The expressed RamR targets a synthetic promoter consisting of BS_RamR_ and UP element, followed by RiboJ-RBS^43, 45^ was implemented to express sfGFP. **c** Fluorescence response of RamR biosensors to NOS. The fluorescence output of SALSOR 0.1 and SALSOR 0.2 in the presence of 0.05% arabinose. WT is *S*. Typhimurium LT2. NOS^+^, 100 μM; NOS^-^, without NOS. To simplify the figure, the RBS upstream of ATF start codon and terminators are not shown. Data are expressed as the mean ± SD of the RFU obtained from three biological replicates. Asterisks indicate a statistically significant difference (*t-*test; ns, not significant; \**p* ≤ 0.05; \*\*\**p* ≤ 0.001). To simplify the figure, the RBS upstream of ATF and terminators are not shown. Abbreviations: AD, activation domain; ATF, artificial transcription factor; a. u., arbitrary units; BS, binding site; JUB1, NAC TF JUNGBRUNNEN1; NOS, noscapine; RBS, ribosome binding site; RNAP, RNA polymerase; RFU, relative fluorescent unit; sfGFP, super-fold green fluorescent protein; TF, transcription factor; TSS, transcription start site; UP, upstream element; WT, wild type. The full data are shown in **Data S4**.

**Fig. 6.**
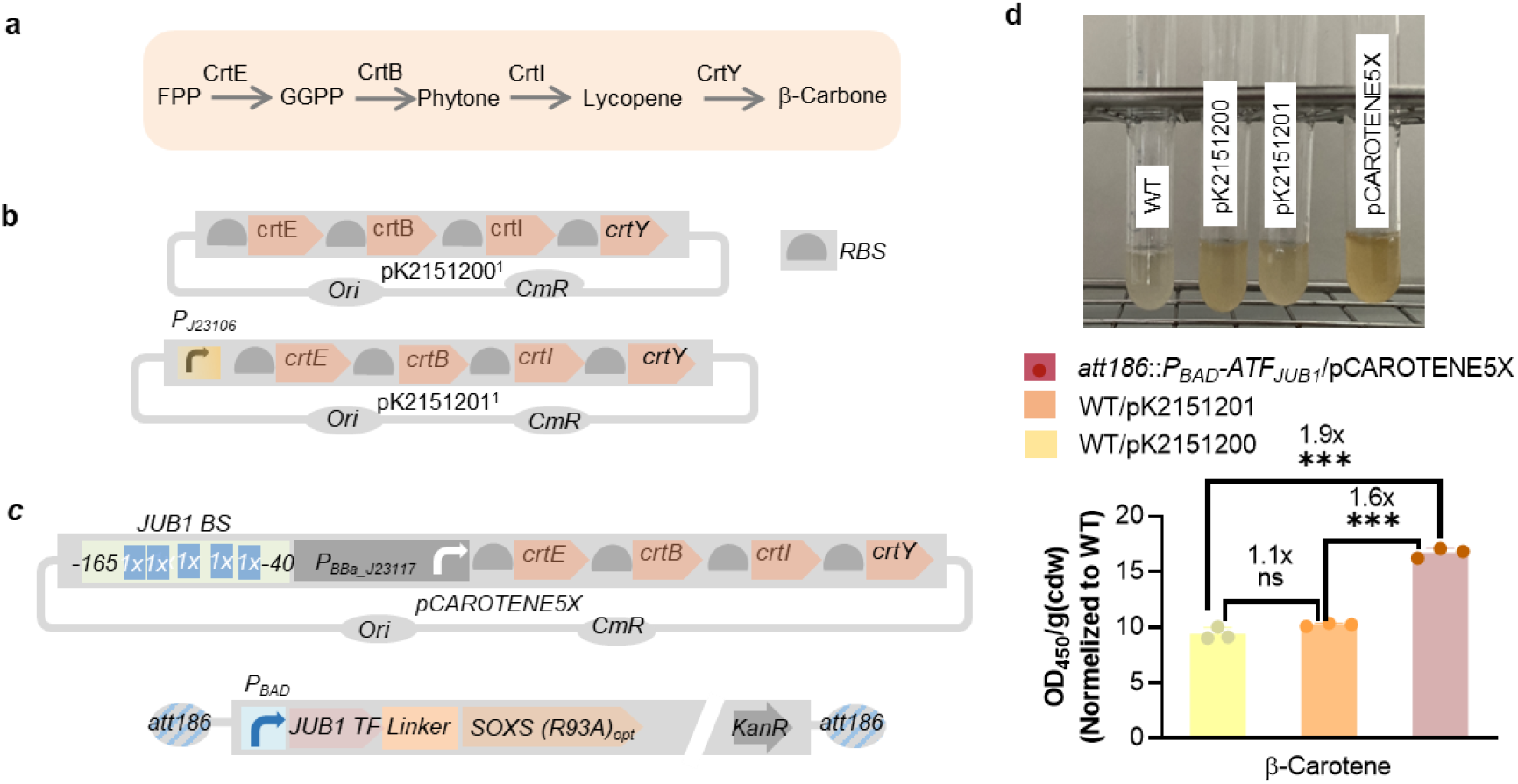
Production of β-carotene in *E. coli* strains. **a** Schematics showing the β-carotene production from bacterial FPP precursor. CrtB, phytoene synthase; CrtY, lycopene cyclase; CrtI, phytoene desaturase; CrtE, geranylgeranyl pyrophosphate synthase. **b** Scheme showing the β-carotene-encoding plasmids used in this study. pK2151200 is results of assembly of BioBricks K118014 (RBS+crtE), K118006 (RBS+crtB), K118005 (RBS+crtI) and K118013 (crtY) in single operon (constructed by Glasgow iGEM 2016). In plasmid pK2151201, bacterial constitutive strong promoter *J23106* (designed by J. Anderson, iGEM2006) controls the expression of the synthetic crtEBIY operon. **c** Scheme representing production of β-carotene using JUB1-derived ATF. In plasmid pCAROTENE5X, *J1*-derivative synthetic promoter containing five copies of plant JUB1 TF, fused to P_BBa_J23117_^24^, controls the expression of “crtEBIY” operon. A JUB1-derived ATF donor contains *P_BAD_*, the full length of JUB1 TF, 5-aa linker^24^, SoxS(R39A)^24^ (that is optimized for expression in *S*. Typhimurium). It additionally encodes KanR. The donor is flanked by 40-bp homology arms to integrate into the *att186* site^39^. To simplify the figure, the RBS upstream of ATF and terminators are not shown. **d** Analysis of carotenoid content of *E. coli* strains. The JUB1-derived ATF donor was integrated into the *att186* site of *E. coli*. Followed by pCAROTEN5X transformation, β-carotene production was quantified in the presence of 0.2% arabinose (WT/pCAROTENE5X). Representative culture of the constructed β-carotene producing strains were shown in top. The β-carotene absorbance at 450 nm was measured using the method reported by Lian *et al*^46^, and was divided to cdw. The data were normalized to that of the WT. Values represent the mean ± SD of three independent colonies in the presence of 0.2% arabinose. *E. coli* strains containing pK2151200 (WT/pK2151200) and pK2151201 (WT/pK2151201) were used as controls. Asterisks indicate a statistically significant difference (*t-*test; ns, not significant; \*\*\**p* ≤ 0.001). “*x*” inside the columns represents the fold induction compared to WT/pK2151200. Abbreviations: aa, amino acid; AD, activation domain; ATF, artificial transcription factor; BS, binding site; cdw, cell dry weight; JUB1, FPP, farnesyl pyrophosphate; GGPP, geranylgeranyl pyrophosphate; NAC TF JUNGBRUNNEN1; KanR, kanamycin resistance marker; RBS, ribosome binding site; TF, transcription factor; WT, wild type. Full data are shown in **Data S5**.

### Arabinose-inducible, plant-derived ATF for β-carotene production in *E. coli*

We next aimed to validate the capacity of plant-derived ATFs to activate transcription of large operons for metabolic engineering applications in Gram-negative bacteria. To this end, we chose to evaluate control of the well-characterized β-carotene biosynthesis pathway using plant-derived ATFs. In order to convert bacterial farnesyl pyrophosphate (FPP) precursor to β-carotene, geranylgeranyl diphosphate synthase (*crtE*), phytoene synthase (*crtB*), phytoene desaturase (*crtI*) and lycopene cyclase *(crtY)* enzymes are needed (**Fig. 6a**). We first transformed *E. coli* with the episomal pK2151201 plasmid (constructed by Glasgow iGEM 2016, **Fig. 6b**) harboring the *crtEBIY* genes in a single operon placed under control of the constitutive, strong promoter *P_J23106_* (designed by J. Anderson, iGEM2006), and where each gene is placed downstream of an individual RBS. We next implemented a synthetic promoter containing five copies of the JUB1 TF binding site to control expression of the *crtEBIY* operon in pCAROTENE5X and integrated the arabinose-inducible JUB1-derived ATF into the *att186* site of *E. coli*. The episomal pK2151200 plasmid harboring the *crtEBIY* operon without a promoter (constructed by Glasgow iGEM 2016, **Fig. 6b**) was used as control.

As shown in **Fig. 6c**, the strain with ATF control modules produced ~1.9-fold more β-carotene than the control strain that harbored a promoter-less *crtEBIY* operon. Importantly, the β-carotene level was ~1.6-fold increased in the strain expressing the JUB1-derived ATF compared to the strain where a strong *P_J23106_* promoter expressed the *crtEBIY* operon, suggesting that the JUB1-derived ATF has the capacity to increase transcription of the *crtEBIY* operon.

## Discussion

Gram-negative bacteria display biotechnological potential to serve as a platform for the production of hard-to-express proteins and might function as drug delivery system exploiting their large repertoire of protein secretion systems. However, a major problem preventing drug-carrying Gram-negative bacteria from being fully exploited in cancer therapy is the toxicity of the drugs to non-tumor tissues, resulting in damage to healthy tissues at the sites of initial localization of the bacteria (mainly the liver and spleen)^47^. Therefore, inducible systems allowing protein expression at the desired time are needed^48^. In addition, many Gram-negative bacteria harbor important virulence factors under the control of complex, poorly understood gene regulatory networks. In order to better understand the regulatory mechanisms of Gram-negative pathogenesis, but also to optimize them for bioengineering applications, species-specific orthogonal synthetic tools are required to enable inducible gene expression with a wide dynamic range. Yet, such tools are largely absent from the methodological toolbox currently used in prokaryotic synthetic biology research^49, 50, 51^.

It is preferable to use synthetic regulatory systems to induce target gene expression. In the past, a few native promoters of Gram-negative bacteria have been used to control expression of target genes. However, native bacterial promoters are under the control of endogenous bacterial TFs, which may interfere with endogenous regulatory networks of the host, making them suboptimal for synthetic biology applications. Furthermore, the most effective production strategy preferably combines a biomass growth phase followed by a protein-of-interest production phase. Using inducible regulators, the expression of regulators and any possible adverse interactions with the genome, metabolites, and proteins of the host that might result in undesirable fitness costs can be delayed until the optimal time of induction. In this study, we evaluated the usage of arabinose as an inducer of gene expression in *Salmonella*. As the availability of arabinose as a nutrient source plays a critical role in modulating *P_BAD_*-controlled gene expression^32^, we first optimized the arabinose catabolic pathways by modifying the endogenous regulatory network. We next established arabinose-inducible, CRISPR/dCas9-derived ATFs to control gene expression in bacteria. We further evaluated several TF families from the plant *A. thaliana* (absent from prokaryotes) for their capacity to induce gene expression in bacteria. We developed and tested 18 and 11 arabinose-inducible ATFs derived from dCas9 and plant TFs in *Salmonella* (shown in **Fig. 3d**, **Fig. 4c** and **d**, and **Fig. S5**) and *E. coli* (shown in **Fig. 4a** and **Supplementary Fig. 3a**), respectively. Here, we dissected the interplay between TF copy number (using plasmid- and chromosomal-based expressions), the number of its target binding sites, in wild-type background or optimized background (through rewiring the endogenous network allowing weak expression or deletion of the arabinose consuming pathway, **Fig. 2**). We observed a broad spectrum of inducible transcriptional outputs that makes our collection of ATFs a suitable choice for synthetic biology applications in *E. coli* and *Salmonella*. We detected a high degree of regulator transferability from *E. coli^24^* (**Supplementary Fig. 3a**) to *Salmonella* (**Fig. 3d**) in case of the strong CRISPR/dCas9-derived ATF in combination with scRNA targeting J105 within a constitutive promoter. In other words, the strongest CRISPR/dCas9-derived ATF led to the highest gene expression, regardless of the host of choice, suggesting that other available regulatory tools for *E. coli*^24, 40^ can be adapted to establish ATFs in *Salmonella*. Here, we also validated the transferability of terminators and promoters from *E. coli* to *Salmonella*, in addition to plant-derived ATFs and CRISPR/dCas9 ATFs from, respectively, plant and *E. coli*. To demonstrate a practical application of plant-derived ATFs, we studied β-carotene production in *E. coli*, where one of our developed ATF/BSs systems was employed to control gene expression of the β-carotene biosynthesis operon. An *E. coli* strain harboring the β-carotene biosynthesis operon under control of the plant-derived ATF, produced β-carotene to ~1.6-fold greater levels compared to a strain that expressed the β-carotene biosynthesis operon from a constitutive promoter. In fact, the metabolic burden raised from the constitutive expression of the pathway enzymes may cause more severe effects compared to the inducible plant-derived ATF expression system, allowing β-carotene production at a specific time point, by addition of arabinose (e.g. in the production phase), opening up a new door for applying plant-derived ATFs in prokaryotic microbial cell factories. We further demonstrated that plant-derived ATFs are a promising tool for establishing *Salmonella* as a sensitive biosensor detecting alkaloid NOS, by developing a biosensor strain termed SALSOR 0.2. The SALSOR 0.2 strain can be used for high-throughput screening of other alkaloid products in chemical engineering projects. The complete microbial biosynthesis of NOS has recently been reported^52^. Therefore, it might be one next valuable anticancer metabolite to be sustainably biomanufactured in microorganisms such as *E. coli, Salmonella*, and yeast. An interesting potential application of SALSOR 0.2 strain will be its application as the whole-cell biosensor for rapid quantification of the extracellular alkaloid concentration produced by other microbial cell factories^42, 53^. In addition to NOS, SALSOR 0.2 might be utilized to detect other therapeutically relevant and commercially available RamR-interacting BIAs such as papaverine, rotundone and glaucine. Additionally, the plant-derived ATFs can be employed to establish other sensitive biosensors, such as CamR from *Pseudomonas putida* that is able to detect bicyclic monoterpenes alkaloids^54^.

In summary, we established a collection of arabinose-inducible ATF for tunable gene expression in *Salmonella* and *E. coli*, which can be potentially adapted for synthetic biology applications also in other Gram-negative bacteria, such as *Pseudomonas aeruginosa*. Another aspect for further improvement is the assessment of other ATFs derived from TFs of either plants or other heterologous organisms in *Salmonella* and *E. coli* Further, our collection of arabinose-inducible ATFs allows for fine-tuning of gene expression via controlling the heterologous protein production and are therefore promising tools for rewiring gene regulatory networks of Gram-negative bacteria.

## Methods

### General

Strains used in this study derive from *S. enterica* serovar Typhimurium strain LT2, *E. coli* DH10β (NEB, Frankfurt am Main, Germany) and *E. coli* DH10B-ALT (*E. coli* DH10B modified to constitutively express *araC*, Addgene, #61151) and are listed in **Table S1**. Plasmids were constructed using the NEBuilder HiFi DNA assembly strategy of New England Biolabs (NEB, Frankfurt am Main, Germany)^55^, or digestion and ligation using T4-DNA ligase (NEB, Frankfurt am Main, Germany). Plasmid and primer sequences are listed in **Tables S2** and **S3,** respectively. PCR amplification of DNA fragments was performed using high-fidelity polymerases: Q5 DNA Polymerase (New England Biolabs, Frankfurt am Main, Germany), Phusion Polymerase (Thermo Fisher Scientific) or PrimeSTAR GXL DNA Polymerase (Takara Bio, Saint-Germain-en-Laye, France) according to the manufacturers’ recommendations. All restriction enzymes were purchased from New England Biolabs (Frankfurt am Main, Germany). Amplified and digested DNA fragments were gel-purified prior to further use. Primers were ordered from IDT (Integrated DNA Technologies Inc., Dessau-Rosslau, Germany) and Sigma-Aldrich (Deisenhofen, Germany). All gBlocks were ordered from IDT (Dessau-Rosslau, Germany). Standard *E. coli* cloning strains were NEB dam^-^/dcm^-^, NEB 5α, or NEB 10β (New England Biolabs) were transformed by heat-shock to propagate constructed plasmids. Strains were grown in Luria-Bertani (LB) medium containing 10 g Tryptone, 5 g Yeast Extract, 5 g NaCl per liter and an appropriate antibiotic(s) (Ampicillin, 100 μg/ml; Chloramphenicol, 50μg/ml; or Kanamycin, 50 μg/ml). The integrity of plasmid constructs was confirmed by sequencing (Microsynth Seqlab, Goettingen, Germany).

Plasmids or integration fragments were amplified by PCR, and, using λ red-mediated homologous recombination standard protocols, twere next integrated into the chromosome of *S*. Typhimurium LT2 (see **Supplementary Method**), or *E. coli* (NEB). Integrations into chromosomal *attB* sites of *S*. Typhimurium LT2 or *E. coli* DH10B-ALT target strains were performed as described by St-Pierre *et al*. (2013)^58^. DNAs were introduced by electroporation into the cells. For selecting chromosomal integrants, we used the appropriate antibiotic(s) (Ampicillin, 50 μg/ml; Chloramphenicol, 15 μg/ml; or/and Kanamycin, 25 μg/ml). When required, an appropriate concentration of arabinose (see **Results**) was added. Single copy integration of each linearized fragment into the target chromosomal site was verified by colony PCR (and by sequencing). Methodologies to construct the plasmids and strains were described in **Supplementary Methods**.

### Induction experiments

Single colonies of bacterial reporter strains were inoculated into 2 mL LB supplemented with appropriate antibiotics and grown at 37 °C (*E. coli*) or 30 °C (*S*. Typhimurium LT2), 220 RPM overnight. Late stationary phase cultures were diluted 1:100 in LB supplemented with appropriate antibiotic(s). For inducible system construction with *P_BAD_*, strains were inoculated in LB medium supplemented appropriate antibiotic(s) with arabinose at 0.0001%, 0.0005%, 0.001%, 0.005%, 0.01%, 0.05%, 0.1%, or 0.2% and cells were harvested at certain time point (see **Results**).

### Plate reader experiments

Cells were inoculated in 2 mL LB supplemented with appropriate antibiotics for fluorescent reporter experiments and grown at 30 °C (*S* Typhimurium), or 37 °C (*E. coli*), 220 RPM overnight. Overnight cultures were diluted 1:100 in fresh LB with appropriate antibiotic(s) (see “**Supplementary Method**”), and appropriate concentration of arabinose (see **Results**) and 150 μL were aliquoted in triplicate flat, clear-bottomed 96-well black plates (Greiner Bio-one), closed with corning 96 well plates sterile lid and condensation rings (Greiner Bio-one), and grown with shaking at 30 °C (*S* Typhimurium), or 37 °C (*E. coli)* in a Biotek Synergy H1 plate reader. OD_600_ and mRFP1 fluorescence (excitation 510 ± 10 nm, emission 625 ± 20 nm), or OD_600_ and sfGFP fluorescence (excitation 478 ± 10 nm, emission 515 ± 20 nm) were measured every 10 min. For ATF library characterization (**Fig. 3c** and **Fig. 4d**), the fluorescence measured after 210 min was divided by the average OD_600_.

For biosensor characterization in *Salmonella* strains, we adapted the protocol reported by d’Oelsnitz *et al* (2022)^43^ for biosensor characterization in *E. coli* The cells were inoculated in 2 mL LB supplemented with appropriate antibiotics and grown at 30 °C, 220 RPM overnight. The following day, 20 μl of each culture was then used to inoculate six separate wells in a 2 mL 24 Deep Well RB Block (Thermo Scientific) closed with a Seal film (Thermo Scientific) containing 900 μl LB medium, one for test alkaloid ligand NOS and a solvent control. After 3.5 h of growth at 30 °C, cultures were induced with 100 μl LB medium containing either 10 μl DMSO or 100 μl LB medium containing the target alkaloid dissolved in 10 μl DMSO. The maximum NOS concentration of 100 μM was used due to the compound’s solubility limit in 1% DMSO. Cultures were grown for an additional 7 h at 37 °C and 250 RPM and subsequently centrifuged (3,500g, 4 °C, 10 min). The supernatant was removed, and cell pellets were resuspended in 1 ml PBS (137 mM NaCl, 2.7 mM KCl, 10 mM Na_2_HPO_4_, 1.8 mM KH_2_PO_4_, pH 7.4). 100 μL of the cell resuspension for each condition was transferred to a 96-well microtiter sterile black, clear bottom plate closed with 96-well lid (Greiner Bio-One) to perform plate reader experiment.

### Flow cytometry and data analysis

For fluorescent reporter experiments, cells were inoculated in 2 mL LB supplemented with appropriate antibiotics and grown at 30 °C, 220 RPM overnight. Overnight cultures were diluted 1:100 in fresh 2 mL LB with appropriate antibiotics (see “*Construction of plasmids and strains*”) and appropriate concentrations of arabinose (see **Results**) and grown at 37 °C (*S* Typhimurium), 220 RPM. After 210 min, cultures were then diluted 1:40 in PBS and analyzed using a SH800S cell sorter Flow Cytometer (Sony). sfGFP fluorescence values were obtained from a minimum of 10,000 cells in each sample. The mean GFP fluorescence per cell was calculated using FlowJo Software.

For optimization of the arabinose-based toolkit in *Salmonella* (**Fig. 2b**), cells were inoculated in 2 mL LB supplemented. Ampicillin was added to the cultures for plasmid-based expression. The cultures were grown at 37 °C, 220 RPM overnight. Overnight cultures were diluted 1:100 in fresh 2 mL LB (Ampicillin for plasmid-based expression was used), and different concentrations of arabinose were added to the cultures and grown at 37 °C, 220 RPM for 150 min (chromosome based expression) or 180 min (plasmid-based expression). Next, the OD_600_ was measured and cells were fixed in a 4% paraformaldehyde (PFA) solution. Briefly, the cultures were centrifugated. The supernatants were discarded, and the pellets were resuspended in 500 μL of 4 % PFA. After incubation for at least 5 min at room temperature, the cells were washed with PBS before the flow cytometry analysis.

### β-Carotene production and quantification

The *E. coli* strains transformed with pK2151200, and pK2151201 plasmids were plated on LB, supplemented with chloramphenicol. The *E. coli* strains genetically modified at the *att186* site for arabinose-dependent induction of JUB1-derived ATF and transformed with pCAROTENE5X plasmid were plated on LB, supplemented with kanamycin and chloramphenicol. Cells were grown at 30 °C for 24 h. The colonies were inoculated into a 4 mL non-induction LB medium and grown overnight at 30°C and 200 RPM in a rotary shaker. A day after, the pre-cultures were used to inoculate main cultures (4 ml) in an induction medium (0.2% arabinose). All cultures were inoculated from pre-cultures to an initial OD_600_ of 0.1. Cells were grown for 24 h at 30°C and 200 RPM to saturation.

Stationary phase bacterial cells were collected by centrifugation at 13,000×g for 1 min and, using the method reported by Lian *et al.,^46^* β-carotene was assessed. Briefly, cell pellets were resuspended in 1 ml of 3 N HCl, boiled for 5 min, and cooled in an ice-bath for 5 min. Next, the lysed cells were washed with ddH_2_O and resuspended in 400 μl acetone to extract β-carotene. The cell debris were removed by centrifugation. The extraction step was repeated until the cell pellet appeared white. The β-carotene containing supernatant was analyzed for its absorbance at 450 nm (A_450_). The production of β-carotene was normalized to the cell density.

## Data availability

The relevant data are available from the corresponding authors upon request.

## Acknowledgments

We thank Sean Colloms (University of Glasgow) for providing the plasmids pK2151200 and pK2151201, and Kelly T. Hughes (University of Utah) for kindly providing the *Salmonella* strains TH437, TH3730 and TH6701. This work was supported in part by the European Research Council (ERC) under the European Union’s Horizon 2020 research and innovation program (grant to M.E.; agreement no. 864971). E.C. (Max Planck Scientific Member) and M.E. (Max Planck Fellow) acknowledge the Max Planck Society for its support. E. C. also thanks the German Research Foundation (support through the Leibniz Prize).

## Author contributions

M.E. and G.N. conceived the study. G.N. designed experiments. H.R. designed arabinose induction experiments. G.N. analyzed the data with contribution from H.R.. G.N. and M.E. wrote the manuscript. E.C. and M.E contributed funding and resources. All authors proof-read and approved the manuscript.

## Supplementary Figures

**Supplementary Figure 1.**
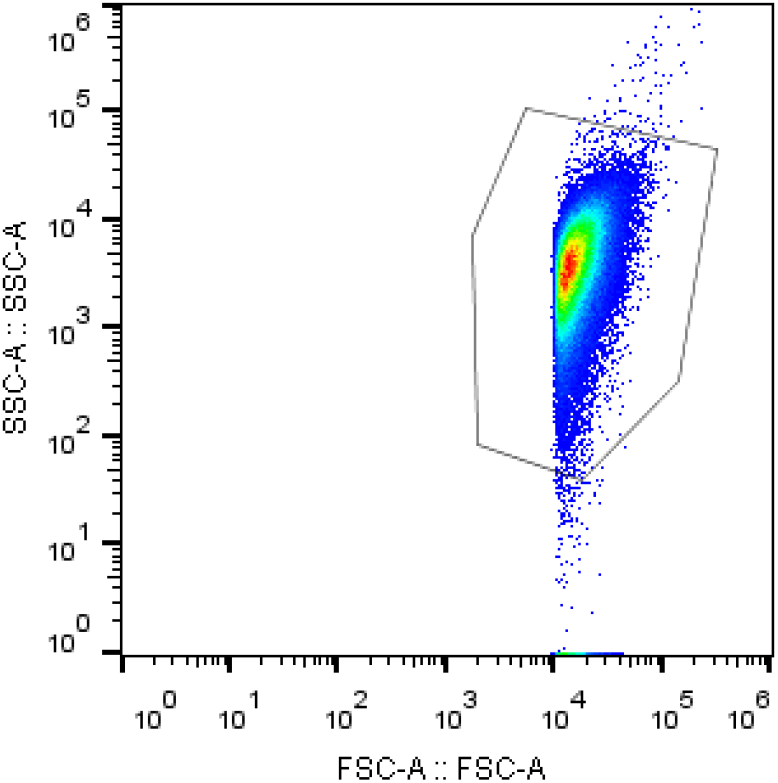
Gating strategy. Bacterial populations were identified by plotting SSC vs. FSC on a log scale. The gated population (in the outlined area) was used for all subsequent measurements of fluorescence. Abbreviations: SSC, side scatter; FSC, forward scatter.

**Supplementary Fig. 2.**
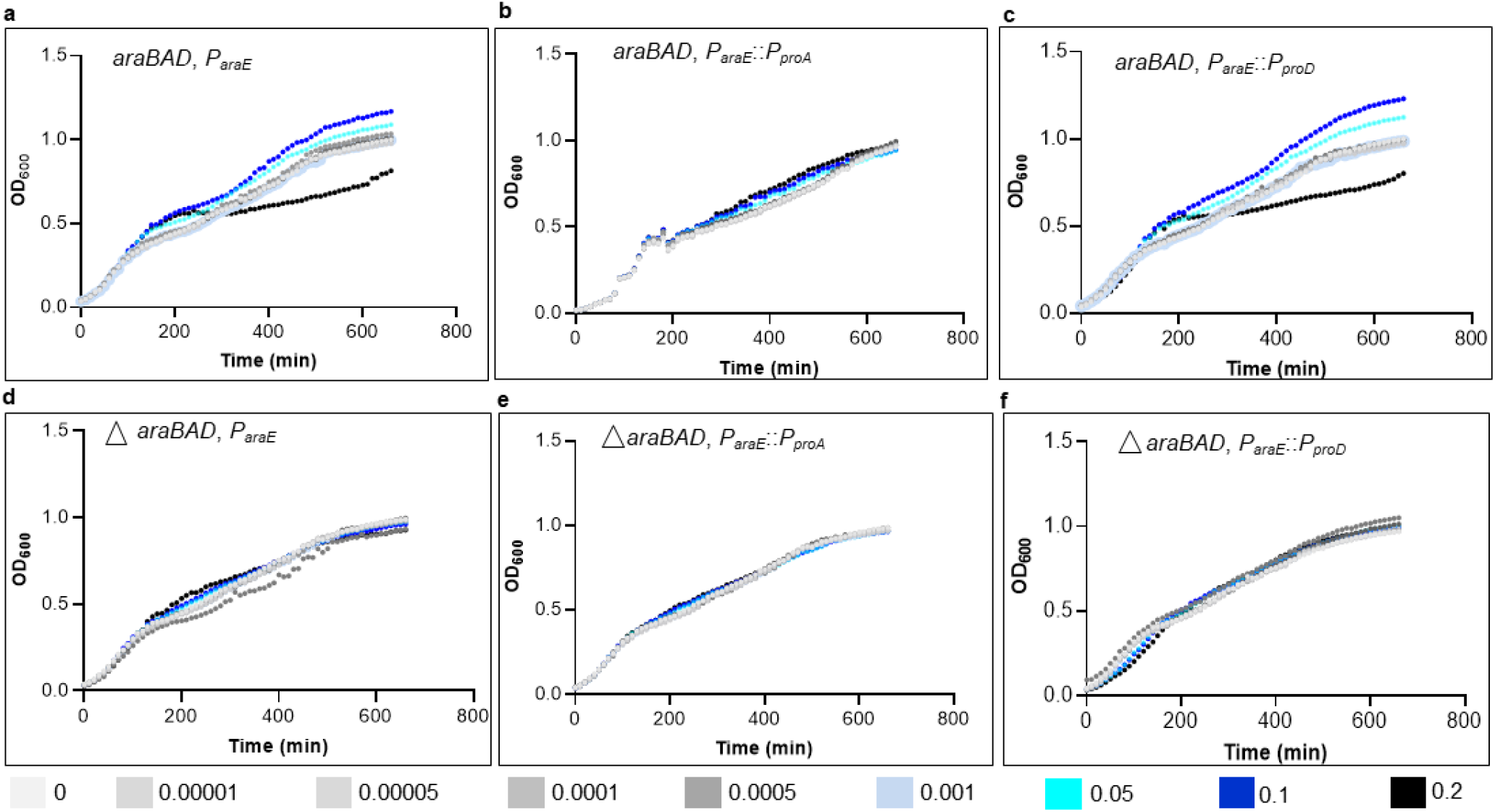
Time-course performance of cells with tuned *araE* expression level treated with various arabinose concentrations. The growth curve of wild-type cells expressing *araE* using **a**) its native promoter *P_araE_*, **b**) arabinose-independent promoters *P_proA_* and **c**) *P_proD_, araBAD*-mutated background, expressing araE using promoters **d**) *P_araE_*, **e**) *P_proA_*, and **f**) *P_proD_* in absence and presence of different arabinose concentrations, including 0.00001%, 0.00005%, 0.0001%, 0005%, 0.001%, 0.05%, 0.1%, and 0.2%. Abbreviations: a.u., arbitrary units; *P_BAD_*, arabinose-responsive promoter; OD_600_, optical intensity at 600 nm. Data are the average obtained from three independent colonies. The full data are shown in **Data S6**.

**Supplementary Fig. 3.**
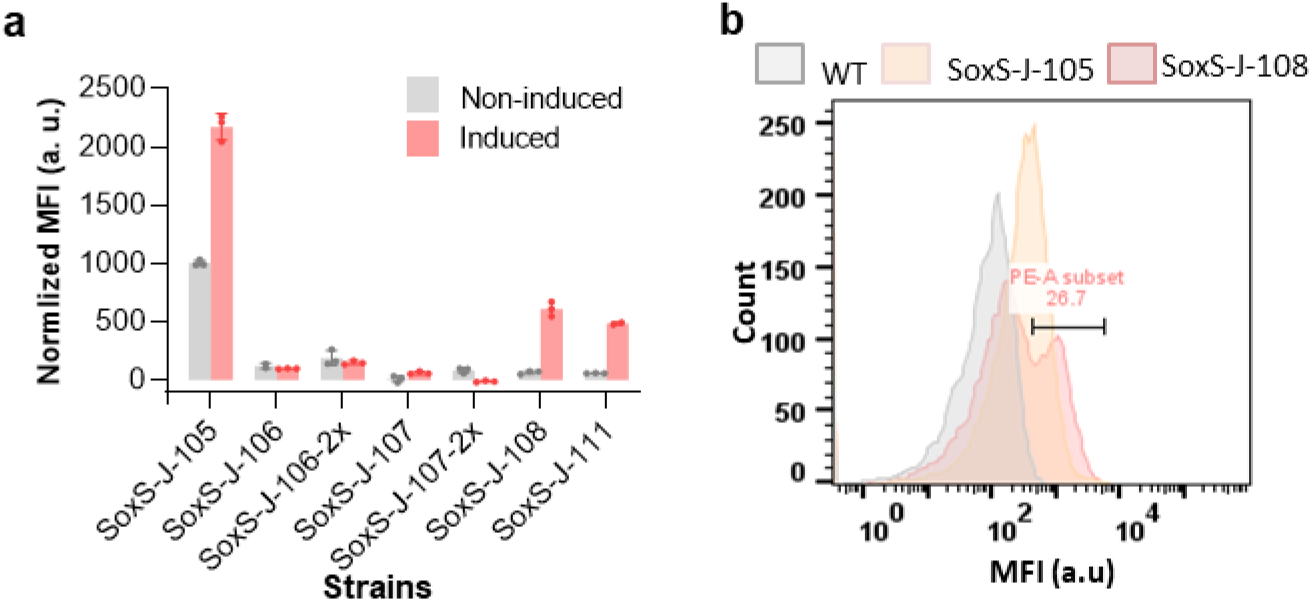
Arabinose-inducible, CRISPR/Cas9-derived ATF for tunable gene expression in *E. coli*. **a** Library of arabinose-inducible, CRISPR/Cas9-derived ATF in *E. coli* 10β. Values reported are mRFP1 fluorescence levels normalized to that of wild-type *E. coli*. Grey, Non-inducing medium, red, inducing medium. Data are expressed as the mean ± SD of the MFI obtained from three independent colonies, normalized to WT *E. coli* 10β. Asterisks indicate a statistically significant difference from the non-inducing medium (*t-*test; \*\*\**p* ≤ 0.001, \*\*\*\**p* ≤ 0.0001). **b** Histogram of mRFP1 fluorescence of *E. coli* 10β cells under the control of CRISPR/Cas9-derived ATF targeting *J105* and J108 in inducing medium. For induction, 0.2% arabinose was used. Abbreviations: a. u., arbitrary units; mRFP1, monomeric red fluorescent protein; MFI, mean fluorescent intensity; WT, wild type. Data are the mean ± SD from three measurements. The full data are shown in **Data S7**.

**Supplementary Fig. 4.**
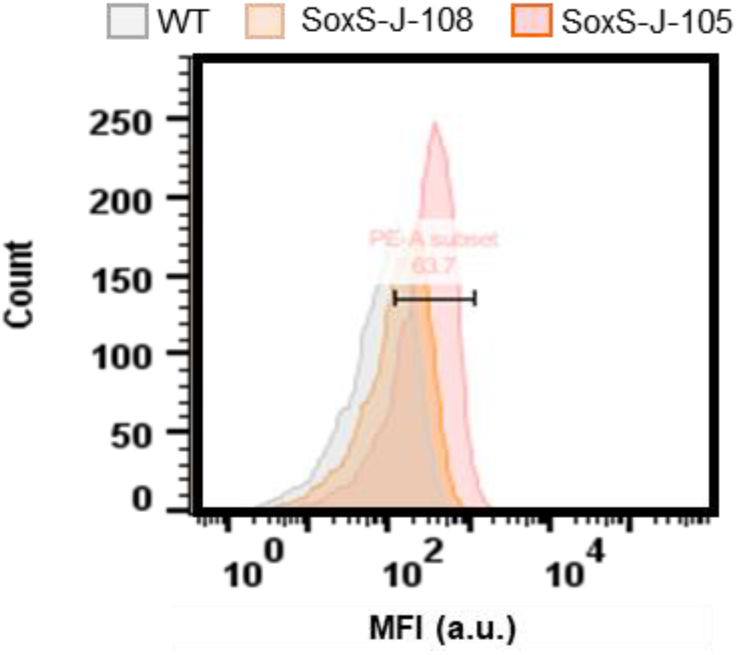
Histogram of mRFP1 fluorescence of *S*. Typhimurium LT2 cells under the control of CRISPR/Cas9-derived ATF targeting *J105* (red) and *J108* (orange) in inducing medium. For induction, 0.05% arabinose was used. Abbreviations: a. u., arbitrary units; mRFP1, monomeric red fluorescent protein; MFI, mean fluorescent intensity; WT, wild type.

**Supplementary Fig. 5.**
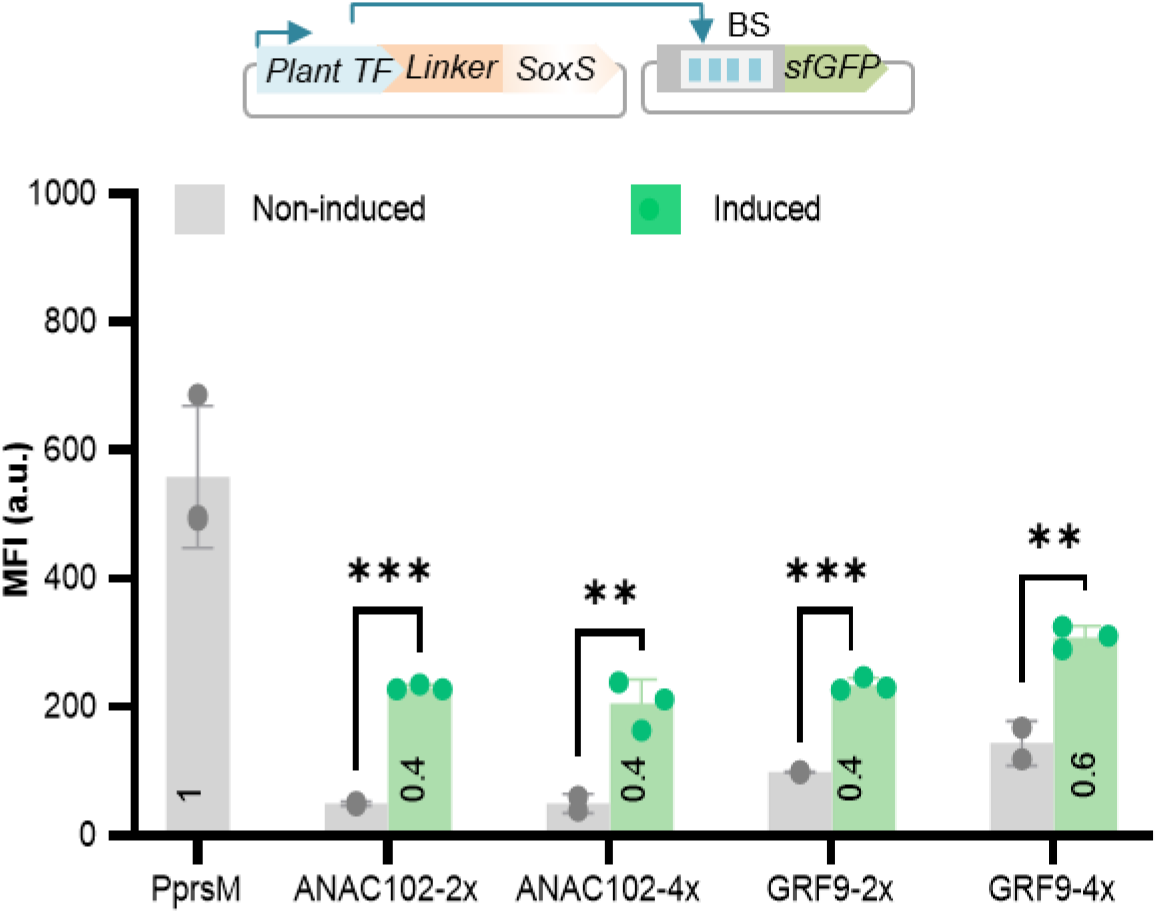
Arabinose-inducible ATFs-derived from plant ANAC102 and GRF9 TFs for *Salmonella*. The transactivation capacity of ANAC102 and GRF9-derived ATFs were tested against the two (2x) and four (4x) copies of its binding site driving sfGFP reporter expression. Fluorescence output is measured in the absence (grey) and presence of 0.05% arabinose (light green). *P_prsM_*, positive control. The MFI of each sample was calculated via FlowJo. Data are expressed as the mean ± SD of the MFI obtained from of three independent colonies. Asterisks indicate a statistically significant difference from the non-inducing medium (*t*-test; ns, not significant; \*\**p* ≤ 0.01; \*\*\**p* ≤ 0.001). Abbreviations: aa, amino acid; AD, activation domain; ATF, artificial transcription factor; a. u., arbitrary units; BS, binding site; ANAC102, NAC TF 102; GRF9, growth regulatory factor; sfGFP, super-fold green fluorescent protein; TF, transcription factor. The full data are shown in **Data S8**.

## References

1. Upadhyaya NM, Ellis JG, Dodds PN. A bacterial Type III secretion-based delivery system for functional assays of fungal effectors in cereals. Humana Press (2014).

2. Widmaier DM, Voigt CA. Quantification of the physiochemical constraints on the export of spider silk proteins by *Salmonella* type III secretion. Microb Cell Fact 9, 78 (2010).

3. Majander K, et al. Extracellular secretion of polypeptides using a modified *Escherichia coli* flagellar secretion apparatus. Nat Biotechnol 23, 475–481 (2005).

4. Nguyen VH, Kim HS, Ha JM, Hong Y, Choy HE, Min JJ. Genetically engineered *Salmonella* Typhimurium as an imageable therapeutic probe for cancer. Cancer Res 70, 18–23 (2010).

5. Forbes NS. Engineering the perfect (bacterial) cancer therapy. Nat Rev Cancer 10, 785–794 (2010).

6. Felgner S, et al. The immunogenic potential of bacterial flagella for *Salmonella*-mediated tumor therapy. Int J Cancer 147, 448–460 (2020).

7. Yoon W, et al. Application of genetically engineered *Salmonella* Typhimurium for interferon-gamma-induced therapy against melanoma. Eur J Cancer 70, 48–61 (2017).

8. Kishimoto H, et al. Tumor-selective, adenoviral-mediated GFP genetic labeling of human cancer in the live mouse reports future recurrence after resection. Cell Cycle 10, 2737–2741 (2011).

9. Pawelek JM, Low KB, Bermudes D. Bacteria as tumour-targeting vectors. Lancet Oncol 4, 548–556 (2003).

10. Hong H, et al. Targeted deletion of the ara operon of *Salmonella* Typhimurium enhances L-arabinose accumulation and drives PBAD-promoted expression of anti-cancer toxins and imaging agents. Cell Cycle 13, 3112–3120 (2014).

11. Oliveira J, Reygaert WC. Gram negative bacteria. In Treasure Island (FL): StatPearls Publishing. PMID: 30855801 (2021).

12. Byndloss MX, Rivera-Chavez F, Tsolis RM, Baumler AJ. How bacterial pathogens use type III and type IV secretion systems to facilitate their transmission. Curr Opin Microbiol 35, 1–7 (2017).

13. Coburn B, Sekirov I, Finlay BB. Type III secretion systems and disease. Clin Microbiol Rev 20, 535–549 (2007).

14. Jahan F, Chinni SV, Samuggam S, Reddy LV, Solayappan M, Su Yin L. The complex mechanism of the *Salmonella typhi* biofilm formation that facilitates pathogenicity: A review. Int J Mol Sci 23 23, 6462 (2022).

15. Husing S, et al. Control of membrane barrier during bacterial type-III protein secretion. Nat Commun 12, 3999 (2021).

16. Macnab RM. How bacteria assemble flagella. Annu Rev Microbiol 57, 77–100 (2003).

17. Pandelakis M, Delgado E, Ebrahimkhani MR. CRISPR-based synthetic transcription factors in vivo: The future of therapeutic cellular programming. Cell Syst 10, 1–14 (2020).

18. Naseri G, Koffas MAG. Application of combinatorial optimization strategies in synthetic biology. Nat Commun 11, 2446 (2020).

19. Davis JH, Rubin AJ, Sauer RT. Design, construction and characterization of a set of insulated bacterial promoters. Nucleic Acids Res 39, 1131–1141 (2011).

20. Cooper KG, Chong A, Starr T, Finn CE, Steele-Mortimer O. Predictable, tunable protein production in *Salmonella* for studying host-pathogen interactions. Front Cell Infect Microbiol 7, 475 (2017).

21. Silva-Rocha R, de Lorenzo V. Mining logic gates in prokaryotic transcriptional regulation networks. FEBS Lett 582, 1237–1244 (2008).

22. Naseri G, Prause K, Hamdo HH, Arenz C. Artificial transcription factors for tuneable gene expression in *Pichia pastoris*. Front Bioeng Biotechnol 9, 676900 (2021).

23. Naseri G, Balazadeh S, Machens F, Kamranfar I, Messerschmidt K, Mueller-Roeber B. Plant-derived transcription factors for orthologous regulation of gene expression in the yeast *Saccharomyces cerevisiae*. ACS Synth Biol 6, 1742–1756 (2017).

24. Dong C, Fontana J, Patel A, Carothers JM, Zalatan JG. Synthetic CRISPR-Cas gene activators for transcriptional reprogramming in bacteria. Nat Commun 9, 2489 (2018).

25. Ho HI, Fang JR, Cheung J, Wang HH. Programmable CRISPR-Cas transcriptional activation in bacteria. Mol Syst Biol 16, e9427 (2020).

26. Bikard D, Jiang W, Samai P, Hochschild A, Zhang F, Marraffini LA. Programmable repression and activation of bacterial gene expression using an engineered CRISPR-Cas system. Nucleic Acids Res 41, 7429–7437 (2013).

27. Liu Y, Wan X, Wang B. Engineered CRISPRa enables programmable eukaryote-like gene activation in bacteria. Nat Commun 10, 3693 (2019).

28. Schilling C, Koffas MAG, Sieber V, Schmid J. Novel prokaryotic CRISPR-Cas12a-based tool for programmable transcriptional activation and repression. ACS Synth Biol 9, 3353–3363 (2020).

29. Naseri G, Behrend J, Rieper L, Mueller-Roeber B. COMPASS for rapid combinatorial optimization of biochemical pathways based on artificial transcription factors. Nat Commun 10 2615 (2019).

30. Fritz G, et al. Single cell kinetics of phenotypic switching in the arabinose utilization system of *E. coli*. PLoS One 9, e89532 (2014).

31. Khlebnikov A, Skaug T, Keasling JD. Modulation of gene expression from the arabinose-inducible araBAD promoter. J Ind Microbiol Biotechnol 29, 34–37 (2002).

32. Vasicek EM, O’Neal L, Parsek MR, Fitch J, White P, Gunn JS. L-arabinose transport and metabolism in *Salmonella* influences biofilm formation. Front Cell Infect Microbiol 11, 698146 (2021).

33. Schleif R. AraC protein, regulation of the l-arabinose operon in *Escherichia coli*, and the light switch mechanism of AraC action. FEMS Microbiol Rev 34, 779–796 (2010).

34. Simon L. D, Ann H. Conversion of the omega-subunit of *Escherichia coli* RNA polymerase into a transcriptional activator or an activation target. Genes & Development 12, 745–754 (1998).

35. Koita K, Rao CV. Identification and analysis of the putative pentose sugar efflux transporters in Escherichia coli. PLoS One 7, e43700 (2012).

36. Jar-How L, Robert J. R, Laurel H, Gary W. Regulation of l-arabinose transport in *Salmonella* Typhimurium LT2. Mol Genet 185, 136 114 (1982).

37. Madar D, Dekel E, Bren A, Alon U. Negative auto-regulation increases the input dynamic-range of the arabinose system of *Escherichia coli*. BMC Syst Biol 5, 111 (2011).

38. Yan Q, Fong SS. Study of in vitro transcriptional binding effects and noise using constitutive promoters combined with UP element sequences in *Escherichia coli*. J Biol Eng 11, 33 (2017).

39. St-Pierre F, Cui L, Priest DG, Endy D, Dodd IB, Shearwin KE. One-step cloning and chromosomal integration of DNA. ACS Synth Biol 2, 537–541 (2013).

40. MacDonald IC, Seamons TR, Emmons JC, Javdan SB, Deans TL. Enhanced regulation of prokaryotic gene expression by a eukaryotic transcriptional activator. Nat Commun 12, 4109 (2021).

41. Yamasaki K, Kigawa T, Seki M, Shinozaki K, Yokoyama S. DNA-binding domains of plant-specific transcription factors: structure, function, and evolution. Trends Plant Sci 18, 267–276 (2013).

42. Zhang J, et al. A microbial supply chain for production of the anti-cancer drug vinblastine. Nature 609, 341–347 (2022).

43. d’Oelsnitz S, et al. Using fungible biosensors to evolve improved alkaloid biosyntheses. Nat Chem Biol 18, 981–989 (2022).

44. Rahmanian-Devin P, Baradaran Rahimi V, Jaafari MR, Golmohammadzadeh S, Sanei-Far Z, Askari VR. Noscapine, an emerging medication for different diseases: A mechanistic review. Evid Based Complement Alternat Med 8402517 (2021).

45. Clifton KP, et al. The genetic insulator RiboJ increases expression of insulated genes. J Biol Eng 12, 23 (2018).

46. Lian J, HamediRad M, Hu S, Zhao H. Combinatorial metabolic engineering using an orthogonal tri-functional CRISPR system. Nat Commun 8, 1688 (2017).

47. Min JJ, et al. Noninvasive real-time imaging of tumors and metastases using tumor-targeting light-emitting Escherichia coli. Mol Imaging Biol 10, 54–61 (2008).

48. Bai F, Li Z, Umezawa A, Terada N, Jin S. Bacterial type III secretion system as a protein delivery tool for a broad range of biomedical applications. Biotechnol Adv 36, 482–493 (2018).

49. Cooper KG, et al. Regulatory protein HilD stimulates *Salmonella* Typhimurium invasiveness by promoting smooth swimming via the methyl-accepting chemotaxis protein McpC. Nat Commun 12, 348 (2021).

50. Hyun J, et al. Engineered Attenuated *Salmonella* Typhimurium expressing neoantigen has anticancer effects. ACS Synth Biol 10, 2478–2487 (2021).

51. Prindle A, et al. Genetic Circuits in *Salmonella* Typhimurium. ACS Synth Biol 1, 458–464 (2012).

52. Li Y, Li S, Thodey K, Trenchard I, Cravens A, Smolke CD. Complete biosynthesis of noscapine and halogenated alkaloids in yeast. Proc Natl Acad Sci U S A 115, 3922–3931 (2018).

53. Siedler S, Stahlhut SG, Malla S, Maury J, Neves AR. Novel biosensors based on flavonoid-responsive transcriptional regulators introduced into *Escherichia coli*. Metab Eng 21, 2–8 (2014).

54. d’Oelsnitz S, Nguyen V, Alper HS, Ellington AD. Evolving a generalist biosensor for bicyclic monoterpenes. ACS Synth Biol 11, 265–272 (2022).

55. Zhang Y, Werling U, Edelmann W. SLiCE: a novel bacterial cell extract-based DNA cloning method. Nucleic Acids Res 40, e55 (2012).

56. Karlinsey JE. λ-Red Genetic Engineering in *Salmonella enterica* serovar Typhimurium. In: Advanced Bacterial Genetics (Use of Transposons and Phage for Genomic Engineering) (2007).

57. Hoffmann S, Schmidt C, Walter S, Bender JK, Gerlach RG. Scarless deletion of up to seven methyl-accepting chemotaxis genes with an optimized method highlights key function of CheM in *Salmonella* Typhimurium. PLoS One 12, e0172630 (2017).

58. St-Pierre F, Cui L, Priest DG, Endy D, Dodd IB, Shearwin KE. One-step cloning and chromosomal integration of DNA. ACS Synth Biol 2, 537–541 (2013).

