## Supplementary Methods for "A regulatory toolkit of arabinose-inducible artificial transcription factors for Gram-negative bacteria"

### Construction of plasmids and strains

*Construction of S.* Typhimurium *araBAD background strains*

EM12441 (*ΔaraBAD*::*sfGFP*): sfGFP together with a downstream terminator and 40-bp overhangs homologous to the regions up- and downstream of the *araBAD* operon was PCR-amplified from the plasmid pEM8317 (lab collection) using primers DaraBAD-sfGFP-rev and DaraBAD-sfGFP-fv. The PCR product was introduced by electroporation into a (S. Typhimurium LT2) strain with a *tetRA* cassette replacing the araBAD operon and harboring the λ-RED-helper plasmid pKD46 (TH6706)[1] to facilitate λ red-mediated replacement of tetRA with sfGFP. Successful recombinants were selected for tetracycline-sensitivity (TcS) using TcS plates[2].

EM12709 (*ΔaraBAD*::*sfGFP*, $P_{araE}$::$P_{proA}$): Strain EM12441 (generated in this study) was transformed with pKD46[1]. In the generated strain EM12472, λ red-mediated recombineering was used to replace $P_{araE}$ first with the *tetRA* expression module. To do this, EM12472 was transformed with PCR-amplified *tetRA* (ParaE-tetRA-fv/ParaE-tetRA-rev on genomic DNA of TH3730, Kelly T. Hughes). Followed by selection on 15 µg/ml tetracycline plate, the generated strain was named EM12498. Next, to subsequently replace *tetRA* with $P_{proA}$, PCR-amplified $P_{proA}$ (primers PproX-araE-fv/PproX-araE-rev on genomic DNA of strain EM9661, lab collection) was transformed into the strain EM12498 to be replaced with t*etRA* (using λ red-mediated recombineering). The generated strain was named EM12709.

EM12746 (*araBAD*, $P_{araE}$::$P_{proA}$): The (native) *araBAD* operon of a *S.* Typhimurium wild type strain (strain TH437, Kelly T. Hughes) was transferred to EM12709 (generated in this study) using bacteriophage $P_{22}$-mediated transduction. To screen the positive clone, selection was performed for capability to use arabinose as the sole carbon source on a carbon-free plate supplemented with arabinose.

EM12886 (*araBAD*/pBAD24-sfGFP): Plasmid pBAD24-sfGFPx1 (Addgene, #51558) was transformed into wild-type *S.* Typhimurium LT2 (TH437, Kelly T. Hughes). To screen the positive clone, selection was performed for capability to use arabinose as the sole carbon source on a carbon-free plate supplemented with arabinose.

EM12877 (*ΔaraBAD*, $P_{araE}$::$P_{proA}$): The $P_{araE}$::$P_{proA}$ fragment of EM12709 (generated in this study) was transferred into TH6701 in which the *araBAD* operon is replaced with *tetRA* (Kelly T. Hughes) using bacteriophage $P_{22}$-mediated transduction.

EM12887 (*ΔaraBAD*/pBAD24-sfGFP): Plasmid pBAD24-sfGFPx1 (Addgene, #51558) was introduced into *S.* Typhimurium LT2, in which the *araBAD* operon is replaced with *tetRA* (TH6701, Kelly T. Hughes).

EM12888 ($P_{araE}$::$P_{proA}$/pBAD24-sfGFP): Plasmid pBAD24-sfGFPx1 (Addgene, #51558) was introduced into strain EM12746 (generated in this study).

EM12889 (*ΔaraBAD*, $P_{araE}$::$P_{proA}$/pBAD24-sfGFP): Plasmid pBAD24-sfGFPx1 (Addgene, #51558) was introduced into strain EM12887 (generated in this study).

EM12967 (*araBAD*, $P_{araE}$::$P_{proD}$): PCR-amplified $P_{proD}$ (primers PproX-araE-fv/PproX-araE-rev on genomic DNA of strain EM8513, lab collection) was introduced into strain EM12498 (generated in this study). Using λ red-mediated recombineering[3], *tetRA* was replaced with $P_{proD}$ to construct strain EM12712. Next, the *araBAD* fragment was transferred from TH437 (Kelly T. Hughes) to EM12712 using $P_{22}$ (Bacteriophage *S.* Typhimurium) transduction to generate the EM12967 strain.

EM13000 (*ΔaraBAD*, $P_{araE}$::$P_{proD}$): *ΔaraBAD925*::*tetRA* fragment of TH6701 was transferred into EM12967, carrying pKD46 (generated in this study) using λ red-mediated recombineering[3].

*Expression plasmids for dCas9-derived ATFs*

pOSIP-KO_PBAD-dCas9: The gBlock "N-$dCas9_{opt}$", containing N-terminal region of dCas9 codon-optimized for expression in *S.* Typhimurium LT2 ($dCas9_{opt}$) was synthesized by IDT (Dessau-Rosslau, Germany). The PCR-amplified "N-$dCas9_{opt}$" fragment (primer pair N-dCas9-fv/N-dCas9-rv, on "N-$dCas9_{opt}$") was cloned in *Asc*I/*Pme*I-digested pL0A0-1-1[4] using the NEBuilder HiFi DNA assembly strategy to generate "pL0A0-1-1_N-dCas9". The gBlock "C-$dCas9_{opt}$-$T_{rrnBT1}$", containing C-terminal of dCas9 that is codon-optimized for expression in *S.* Typhimurium ($dCas9_{opt}$), fused to bacterial *rrnBT1* terminator ($T_{rrnBT1}$), was synthesized by IDT (Dessau-Rosslau, Germany). The PCR-amplified "C-$dCas9_{opt}$-$T_{rrnBT1}$" fragment (primer pair C-dCas9-fv/C-dCas9-rv on gBlock "C-$dCas9_{opt}$-$T_{rrnBT1}$") was cloned into *Xba*I/*Pme*I-digested pL0A0-1-1_N-dCas9 using the NEBuilder HiFi DNA assembly strategy to generate pL0A0-1-1_dCas9. PCR-amplified $P_{BAD}$ fused to a synthetic strong RBS (as described by Dong *et al*. (2018)[5] (primer LacI-AraC-ParaB-fv/LacI-AraC-ParaB-rv on "placI-araC-PBAD-RBS", ATG Bio-Synthesis) was cloned into *Sal*I-digested pL0A-0-1-1_dCas9 using the NEBuilder HiFi DNA assembly strategy to generate "pL0A-1-1_PBAD_dCas9". Next, the PCR-amplified "$P_{BAD}$-dCas9-$T_{rrnBT1}$" fragment (primer pair dCas9-fv/dCas9-rv on "pL0A-1-1_PBAD_dCas9") was cloned into *Bam*HI/*Spe*I-digested pOSIP-KO[6] to generate the "pOSIP-KO_PBAD-dCas9" plasmid.

*Reporter plasmids for dCas9-derived ATFs*

pJ-105: A 1094-bp fragment containing crRNA to target the J106 motif within the J1 promoter, together with its leader and terminator sequences was PCR-amplified using primer pair BBa_J23119-gRNA-J106-fv/BBa_J23119-gRNA-J106-rv from pCK005.6 (Addgene, #153025)[5]. Subsequently, it was cloned into a PCR-amplified 1869 bp fragment of pUC19 (NEB, primer pair PUC19-fv/PUC19-rv) using the NEBuilder HiFi DNA assembly strategy to generate plasmid p106. Next, the J106-targeting crRNA region of plasmid p106 was replaced with crRNA to target the J105 motif within the *J1* region of the synthetic promoter[5]. To this end, the "p105" fragment was PCR-amplified (primer pair crRNA-J105-fv/crRNA-J105-rev on p106) and followed by NEBuilder HiFi DNA assembly-mediated circularisation of the PCR product, the plasmid p105 was generated. Subsequently, p105 was digested with *Acl*I and *Aat*II to remove a 690 bp fragment. The *J1* region of the synthetic promoter fused to mRFP1 and its downstream terminator was PCR-amplified (primer pair J1-RFP-fv/J1-RFP-rv on pJF076Sa, Addgene, #113322[5]) and cloned into the linearized p105 plasmid using the NEBuilder HiFi DNA assembly strategy. The generated plasmid was called "pJ-105".

pSoxS-J-105: PCR-amplified $P_{tetR}$-tetR-$T_{tetR}$ fragment (primer pair TET-R-fv/TET-R-rv on genomic DNA TH3730, gift from Kelly T. Hughes) and PCR-amplified fragment containing coding regions for $P_{BAD}$, MCP optimized for expression in *S.* Typhimurium LT2 (optMCP), a six amino acid linker, SoxS activation domain optimized for expression in *S.* Typhimurium LT2 with enhanced transcriptional activity (SoxS(R93A)$_{opt}$), and synthetic $T_{BBa\_B002}$ terminator (primer pair MCP-SOX-fv/MCP-SOX-rv on gBlock "pAra_optMCP_linker_optSOXA_R93A_BBa_B002")[5] were assembled in *Asc*I/*Bam*HI-digested AVA plasmid[7] using the NEBuilder HiFi DNA assembly strategy. The generated plasmid was named "AVA_TetR_SOXS_a". Next, fragment "*$P_{BAD}$-optMCP-linker-optSOXS_R93A-BBa_$T_{BBa\text{-}B002}$*" (primer pair TET-MCP-SOX-fv /TET-MCP-SOX-rv on "AVA_TetR_SoxS_a") was assembled in PCR-amplified backbone pJ-105 (primer pair J-crRNA-fv/ J-crRNA-rv) to generate plasmid pSoxS-J-105.

pSoxS-J-106: Fragment "*$P_{BAD}$-optMCP-linker-optSOXS_R93A-BBa_$T_{BBa\text{-}B002}$*"(primer pair TET-MCP-SOX-fv /TET-MCP-SOX-rv on "AVA_TetR_SoxS_a") was assembled in PCR-amplified backbone pCK005.6[5] (primer pair J-crRNA-fv/ J-crRNA-rv) to generate plasmid pSoxS-106. Plasmid pSoxS-106 was generated and was digested with *Acl*I and *Aat*II to remove 690 bp. The

J1-containing promoter fused to mRFP1, together with its downstream terminator was PCR-amplified (primer pair J1-RFP-fv/J1-RFP-rv on pJF076Sa, Addgene, #113322)[5] and cloned into the linearized pS-106 plasmid using the NEBuilder HiFi DNA assembly strategy. The generated plasmid was called pSoxS-J-106.

pSoxS-J-106_2x: A 2337-bp fragment harboring a modified *J1* region with two copies of the J-106 motif (see **Results** Section), mRFP1, and bacterial $T_{rrnBT1}$ terminator (J1-RFP-2x-fv/J1-RFP-rv on “pUC57_106_2x”, BioCat Inc., Heidelberg, Germany) were cloned into *Acl*I/*Aat*II-digested pSoxS-106 using the NEBuilder HiFi DNA assembly strategy to construct “pSoxS-J-106-2x” plasmid.

pSoxS-J-107: The J106-targeting crRNA region of plasmid p106 was replaced with crRNA to target J107 motif within the *J1*-containing promoter[5] using the NEBuilder HiFi DNA assembly of PCR-amplified fragment (primer pair crRNA-J107-fv/crRNA-J107-rev on p106) to construct p107. Fragment “*$P_{BAD}$-optMCP-linker-optSOXS_R93A-BBa_$T_{BBa-B002}$*“(primer pair TET-MCP-SOX-fv/TET-MCP-SOX-rv on “AVA_TetR_SOXS_a”) was assembled in PCR-amplified backbone p107 (primer pair J-crRNA-fv/ J-crRNA-rv) to generate plasmid pSoxS-107. Plasmid pSoxS-107 was generated and was digested with *Acl*I and *Aat*II to remove 690 bp. The J1 region of the synthetic promoter fused to mRFP1, together with its downstream terminator was PCR-amplified (primer pair J1-RFP-fv/J1-RFP-rv on pJF076Sa, Addgene, #113322)[5] and cloned into the linearized pS-107 plasmid using the NEBuilder HiFi DNA assembly strategy. The generated plasmid was called pSoxS-J-107.

pSoxS-J-107_2x: A 2337-bp fragment harboring modified *J1* region of the synthetic promoter with two copies of J-107 motif (see **Results** Section), mRFP1 and bacterial $T_{rrnBT1}$ terminator (J1-RFP-2x-fv/J1-RFP-rv on “pUC57_107_2x”, BioCat Inc., Heidelberg, Germany) were cloned into *Acl*I/*Aat*II-digested pSoxS--106 using the NEBuilder HiFi DNA assembly strategy to construct “pSJ-107-2x” plasmid.

pSoxS-J-108: The J106-targeting crRNA region of plasmid p106 was replaced with crRNA to target J108 motif within J1 promoter[5] using the NEBuilder HiFi DNA assembly of PCR-amplified fragment (primer pair crRNA-J108-fv/crRNA-J108-rev on p106) to construct p108. Fragment “*$P_{BAD}$-optMCP-linker-optSOXS_R93A-BBa_$T_{BBa-B002}$”* (primer pair TET-MCP-SOX-fv/TET-MCP-SOX-rv on “AVA_TetR_SoxS_a”) was assembled in PCR-amplified backbone p108 (primer pair J-crRNA-fv/ J-crRNA-rv) to generate plasmid pSoxS--108. Plasmid pS-108 was generated and was digested with *Acl*I and *Aat*II to remove 690 bp. The *J1* region of synthetic promoter fused to mRFP1, and its downstream terminator was PCR-amplified (primer pair J1-RFP-fv/J1-RFP-rv on pJF076Sa, Addgene, #113322[5]) and cloned into the linearized pSoxS--108 plasmid using the NEBuilder HiFi DNA assembly strategy. The generated plasmid was called pSJ-108.

pSJ-111: The J106-targeting crRNA region of plasmid p106 was replaced with crRNA to target J111 motif within the *J1* region of synthetic promoter[5] using the NEBuilder HiFi DNA assembly of PCR-amplified fragment (primer pair crRNA-J111-fv/crRNA-J111-rev on p106) to construct p111. Fragment "*$P_{BAD}$-optMCP-linker-optSOXS_R93A-BBa_$T_{BBa\text{-}B002}$*" (primer pair TET-MCP-SOX-fv/TET-MCP-SOX-rv on "AVA_TetR_SOXS_a") was assembled in PCR-amplified backbone p108 (primer pair J-crRNA-fv/ J-crRNA-rv) to generate plasmid pSoxS--111. Plasmid pS-108 was generated and was digested with *Acl*I and *Aat*II to remove 690 bp. J1 promoter fused to mRFP1, and its downstream terminator was PCR-amplified (primer pair J1-RFP-fv/J1-RFP-rv on pJF076Sa, Addgene, #113322[5]) and cloned into the linearized pS-111 plasmid using the NEBuilder HiFi DNA assembly strategy. The generated plasmid was called "pSoxS-J-111".

*dCas9 driver/reporter strains*

"E. coli_att186::PBAD-dCas9": *E. coli* strain DH10B-ALT (Addgene, #61151), constitutively expressing *araC*, was used as the target strain for integration of the dCas9 cassette. The plasmid "pOSIP-KO_PBAD-dCas9" was introduced into the cells, and $P_{BAD}$-dCas9 cassette was clonetegrated into the *att186* site[6]. The positive clones were selected on LB supplemented with Kanamycin (25 μg/ml).

"E_SoxS-J-105", "E_SoxS-J-106", "E_SoxS-J-106_2x", "E_SoxS-J-107", "E_SoxS-J-107_2x", "E_pSoxS-J_108" and "E_SoxS-J_111": Strain "E. coli, att186::PBAD-dCas9" was transformed with reporter plasmids pSoxS-J-105, pSoxS-J-106, pSoxS-J-106_2x, pSoxS-J-107, pSoxS-J-107_2x, pSoxS-J_108 and pSoxS-J_111. The positive clones were grown on LB supplemented with Kanamycin (25 μg/ml) and Ampicillin (50 μg/ml).

"LT2, att186::PBAD-dCas9": *Salmonella* strain TH437 (Kelly T. Hughes), natively expressing *araC*, was used as the target strain for integrating the dCas9 cassette. The plasmid pOSIP-KO_PBAD-dCas9 was introduced into the cells, and PBAD-dCas9 cassette was clonetegrated allowing integration into the *att186* site[6]. The positive clones were selected on LB supplemented with Kanamycin (25 μg/ml).

"LT2_J-105", "LT2_SoxS-J-105, LT2_no gRNA", "LT2_SoxS-J-106", "LT2_SoxS-J-106_2x", "LT2_SoxS-J-107", "LT2_SoxS-J-107_2x", "LT2_SoxS-J-108" and "LT2_SoxS-J-111": Strain "*LT2*, att186::PBAD-dCas9" was transformed with reporter plasmids pJ-105, pSoxS-J-105, pJF076Sa, pSoxS-J-106, pSoxS-J-106_2x, pSoxS-J-107, pSoxS-J-107_2x, pSoxS-J-108 and pSoxS-J-111 to generate strains. The positive clones were grown on LB supplemented with Kanamycin (25 μg/ml) and Ampicillin (50 μg/ml).

*Plant-derived ATF and promoter pair clones*

pJUB1, pJUB1DBD, pANAC102 and pGRF9: PCR-amplified JUB1 full-length (primer pair JUB-fv/JUB-rv, on "pGN003B-JUB1"), JUB1 DBD (primer pair JUBDBD-fv/JUB-rv, on "pGNPP004"), ANAC102 full-length (primer pair ANAC-fv/JUB-rv, on "pGNPP0033") or GRF9 full-length (primer pair GRF-fv/JUB-rv, on "pGNPP002") was assembled in PCR-amplified backbone plasmid containing the 5-aa linker fused to SoxS(R93A) activation domain optimized for expression in *S.* Typhimurium ($SoxS(R93A)_{opt}$) allowing for the arabinose-controlled expression of ATF (primer pair PBAD-SOX-fv/PBAD-SOX-rv, on "AVA_TetR_SOXS_a", see "**Reporter plasmids for dCas9-derived ATFs**") using the NEBuilder HiFi DNA assembly strategy. The generated plasmids were pJUB1, pJUBDBD, pANAC102 and pGRF9.

pJUB2X: a synthetic fragment containing *Sal*I recognition site, two copies of JUB1 binding site fused to 3'-end of *J1* region of the synthetic promoter (84 bp upstream of ATG) and 171-bp of 5'-end of sfGFP ended with *Nco*I site at 3' was designed and generated by BioCat Inc. (Heidelberg, Germany), and cloned in plasmid pUC57 (pUC57-BsaI-free_ JUB2x). The plasmid was digested with *Sal*I and *Nco*I to obtain a 359-bp fragment. Next, the 359-bp fragment was cloned in *Sal*I/*Nco*I-digested pKH70-PrpsM-sfGFP (2632 bp in length, lab collection) using T4 ligase (NEB). The generated plasmid was called pJUB2X.

pJUB0X: Annealed oligonucleotide JUB0X-fv/JUB0X-rv was assembled in *Sal*I/*Xcm*aI-digested pJUB2X to remove JUB1 binding sites. The generated plasmid was called pJUB0X.

pJUB1X, pJUB4X, pANAC2X, pANAC4X, pGRF2X and pGRF4X: The plasmid pJUB2X was digested with *Sal*I/*Xcm*I to remove 186 bp. One, two or four copies of the binding site of JUB1, ANAC102 and GRF9 were PCR-amplified from, respectively, "pGN005B-1xBS-JUB1"[7], "pGN005B-4xBS-JUB1"[7], "pGN005B-2xBS-JANAC102"[7], "pGN005B-4xBS-ANAC102"[7], "pGN005B-2xBS-GRF9"[7] and "pGN005B-4xBS-GRF9"[7] (primer pair BS-fv/BS-rv), and assembled in linearized pJUB2X plasmid using the NEBuilder HiFi DNA assembly strategy to generate "pJUB1X", "pJUB4X", "pANAC2X", "pANAC4X", "pGRF2X" and "pGRF4X".

pJUB5X: A linearized DNA harboring five copies of the JUB1 binding site was PCR-amplified from plasmid pJUB4X using primer pair JUB5X-fv/JUB5X-rv, and circular plasmid pJUB5X was generated via the NEBuilder HiFi DNA assembly of the linearized DNA.

pJUB1-6x-sfGFP: A linearized DNA harboring six copies of the JUB1 binding site was PCR-amplified from plasmid pJUB5X using primer pair JUB5X-fv/JUB5X-rv, and circular plasmid “pJUB6X” was generated via the NEBuilder HiFi DNA assembly of the linearized DNA.

“pKO_PBAD-JUB1” and “pKO-PBAD-JUB1DBD”: PCR-amplified “$P_{BAD}$-JUB1-linker-SOXS(R93A)$_{opt}$-$T_{rrnBT1}$” (primer pair plantTF-fv/plantTF-rv on pJUB1) or “$P_{BAD}$-DBD of JUB1-linker- SOXS(R93A)$_{opt}$-$T_{rrnBT1}$” (primer pair plantTF-fv/plantTF-rv on “pJUB1DBD”) was cloned into *Bam*HI/*Spe*I-digested pOSIP-KO[6] to generate plasmids “pOSIP-KO_PBAD-JUB1” and “pOSIP-KO_PBAD-JUB1DBD”, respectively.

*Plant driver/reporter strains*

"*E. coli*_pJUB1" and "*E. coli*_pJUB1DBD": pJUB1 and pJUB1DBD were introduced into DH10B-ALT via heat-shock method. The positive clones were grown on LB supplemented with Kanamycin (25 μg/ml).

“*E. coli*_pJUB1_pJUB2X” and “*E. coli*_pJUB1DBD_pJUB2X”: “*E. coli*_pJUB1” and “*E. coli*_pJUB1DBD” were transformed with pJUB2X. The positive clones were grown on LB supplemented with Kanamycin (25 μg/ml) and Ampicillin (50 μg/ml).

"*E. coli*_ att186:: PBAD-JUB1" and "*E. coli*_ att186:: PBAD-JUB1DBD": *E. coli* strain DH10B-ALT (Addgene, #61151), constitutively expressing araC, was used as the target strain for integration of JUB-derived ATF cassettes. The plasmid “pKO_PBAD-JUB1” or “pKO-PBAD-JUB1DBD” was introduced into the cells, allowing integration into the *att186* site was clonetegrated using a kanamycin-resistant pOSIP[6]. The positive clones were grown on LB supplemented with Kanamycin (25 μg/ml).

"*E. coli*_ att186:: PBAD-JUB1_pJUB2X" and "*E. coli*_att186:: PBAD-JUB1DBD_pJUB2X ": "*E. coli*_att186:: PBAD-JUB1" and "*E. coli*_att186:: PBAD-JUB1DBD" were transformed with reporter plasmids pJUB2X. The positive clones were grown on LB supplemented with Kanamycin (25 μg/ml) and Ampicillin (50 μg/ml).

"LT2_pJUB1": pJUB1 was introduced into *S.* Typhimurium LT2 via electroporation. The positive clones were grown on LB supplemented with Kanamycin (25 μg/ml).

“LT2_pJUB1_pJUB0X”, “LT2_pJUB1_pJUB1X”, “LT2_pJUB1_pJUB4X”, “LT2_pJUB1_ pJUB5X” and “LT2_pJUB1_pJUB6X”: Strain “LT2_pJUB1” was transformed with plasmids pJUB0X, pJUB1X, pJUB2X, pJUB4X, pJUB5X or pJUB6X. The positive clones were grown on LB supplemented with Kanamycin (25 μg/ml) and Ampicillin (50 μg/ml).

"LT2_PRSM": Plasmid pKH70-PrpsM-sfGFP (lab collection) was introduced into *S.* Typhimurium LT2 via electroporation. The positive clones were grown on LB supplemented with Ampicillin (50 μg/ml).

“LT2_pANAC102_pANAC2X” and “LT2_pANAC102_pANAC4X”: pANAC102 was introduced into *S.* Typhimurium LT2 via electroporation to generate strain “LT2_ pANAC102”. The positive clones were grown on LB supplemented with Kanamycin (25 μg/ml). Plasmids “pANAC2X” and “pANAC4X” were introduced into strain “LT2_pANAC102” to generate strains “LT2_pANAC102_pANAC2X” and “LT2_pANAC102_pANAC4X”, respectively. The positive clones were grown on LB supplemented with Kanamycin (25 μg/ml) and Ampicillin (50 μg/ml).

"LT2_pGRF9_pGRF2X" and "LT2_pGRF9_pGRF4X": pGRF9 was introduced into *S.* Typhimurium LT2 via electroporation to generate strain "LT2, pGRF9". The positive clones were grown on LB supplemented with Kanamycin (25 μg/ml). Plasmids "pGRF2X" and "pGRF4X" were introduced into strain "LT2_pGRF9" to generate strains "LT2_pGRF9_pGRF2X" and "LT2_pGRF9_pGRF4X", respectively. The positive clones were grown on LB supplemented with Kanamycin (25 μg/ml) and Ampicillin (50 μg/ml).

“LT2_ΔaraBAD_PproA::ParaE_pJUB1_pJUB2X” and “LT2_ΔaraBAD_PproA::ParaE_pJUB1_pJUB4X”: Plasmid pJUB1 was introduced into strain “LT2_ΔaraBAD_PproA::ParaE”. The positive clones were grown on LB supplemented with Kanamycin (25 μg/ml). Next, plasmids pJUB2X and pJUB4X were introduced into strain “LT2_ΔaraBAD_PproA::ParaE_pJUB1” to generate strains “LT2_ΔaraBAD_PproA::ParaE_pJUB1_pJUB2X” and “LT2_ΔaraBAD_PproA::ParaE_JUB1_pJUB4X”, respectively. The positive clones were grown on LB supplemented with Kanamycin (25 μg/ml) and Ampicillin (50 μg/ml).

*Biosensor strain*

SALSOR 0.1:

To establish alkaloid biosensor strain[8], a 136-bp fragment between RamR (STM580) and its responsive promoter , controlling RamA (STM581) expression in *S.* Typhimurium LT2 (NC003197, gift from Kelly T. Hughes ) was replaced with PCR-amplified *KanSceI* cassette (primer pair RamR-Kan-fv/RamR-Kan-rv, on pWRG717[3]) using λ red-mediated homologous recombination standard protocols. The generated strain was called RKLT2. The "SELEX alkaloid sensor" containing the native promoter of *ramR* and its RBS (to deliver RamR expression) and RamR-responsive operator from the promoter of *ramA* (to be targeted by RamR) fused to synthetic minimal promoter region (UP element and RiboJ RBS), sfGFP and rrnBT1 terminator was designed and synthesized by ATG bioscience. Next, the "KanSceI" cassette of RKLT2 was replaced with a PCR-amplified "SELEX alkaloid sensor" cassette (primer pair RamR-Sensor-fv/RamR-Sensor-rv, on "SELEX alkaloid sensor") using λ-RED-mediated homologous recombination standard protocols. The generated strain was called SALSOR 0.1.

SALSOR 0.2: The "KanSceI" cassette of RKLT2 was replaced with the "ATF alkaloid sensor" cassette, containing a synthetic promoter with four copies of JUB1 binding site and synthetic RBS (to deliver RamR expression) and RamR-responsive operator from the promoter of *ramA* (to be targeted by RamR) fused to synthetic minimal promoter region (UP element and RiboJ RBS), sfGFP, and rrnBT1 terminator was PCR-amplified from synthetic fragment synthesized by ATG bioscience (primer pair RamR-Sensor-fv/RamR-Sensor-rv) using λ red-mediated homologous recombination standard protocols. The generated strain was called SALSOR4X. Next, plasmid pJUB1 was introduced into SALSOR4X via electroporation to generate SALSOR 0.2 strain.

*β-Carotene-encoding plasmids*

pCAROTENE5X: The origin of replication, Ampicillin resistance gene and synthetic promoter harboring five copies of JUB1 BS were PCR-amplified from pJUB1-5X ("**Plant-derived ATF and promoter pair clones** ") using primer pair 5X-fv and 5X-rv. Fragments K118014(RBS+crtE), K118006(RBS+crtB), K118005(RBS+crtI) and K118013(crtY) were PCR-amplified from pK2151200 (BBa_K2151200 assembled into pSB1C3, gifted by S. Colloms) using primer pair K2151200-fv and K2151200-rv. The two PCR fragments were assembled using the NEBuilder HiFi DNA assembly strategy to generate "pCAROTENE5X".

β-Carotene-producing strain

*E. coli*_pK2151200: Plasmids pK2151200 was introduced into *E. coli* strain 10β (NEB) via the heat shock method.

*E. coli*_pK2151201: Plasmids pK2151201 was introduced into *E. coli* strain 10β (NEB) via the heat shock method.

*E. coli*, att186:: PBAD-JUB1_pCAROTENE5X: plasmid pCAROTENE5X was introduced into *E. coli*, att186:: PBAD-JUB1) via the heat shock method.

### References


1. Datsenko KA, Wanner BL. One-step inactivation of chromosomal genes in *Escherichia coli* K-12 using PCR products. *ACS Synth Biol* **97**, 6640-6645 (2000).

2. Bochner BR, Huang HC, Schieven GL, Ames BN. Positive selection for loss of tetracycline resistance *J Bacteriol* **143**, 926 (1980).

3. Hoffmann S, Schmidt C, Walter S, Bender JK, Gerlach RG. Scarless deletion of up to seven methyl-accepting chemotaxis genes with an optimized method highlights key function of CheM in *Salmonella* Typhimurium. *PLoS One* **12**, e0172630 (2017).

4. Naseri G, Behrend J, Rieper L, Mueller-Roeber B. COMPASS for rapid combinatorial optimization of biochemical pathways based on artificial transcription factors. *Nat Commun* **10**, 2615 (2019).

5. Dong C, Fontana J, Patel A, Carothers JM, Zalatan JG. Synthetic CRISPR-Cas gene activators for transcriptional reprogramming in bacteria. *Nat Commun* **9**, 2489 (2018).

6. St-Pierre F, Cui L, Priest DG, Endy D, Dodd IB, Shearwin KE. One-step cloning and chromosomal integration of DNA. *ACS Synthetic Biology* **2**, 537-541 (2013).

7. Naseri G, Prause K, Hamdo HH, Arenz C. Artificial transcription factors for tuneable gene expression in *Pichia pastoris*. *Front Bioeng Biotechnol* **9**, 676900 (2021).

8. d'Oelsnitz S*, et al.* Using fungible biosensors to evolve improved alkaloid biosyntheses. *Nat Chem Biol*, 1038 (2022).
